# A developmental condensin I complex assists the *Paramecium* PiggyMac domesticated transposase during programmed DNA elimination

**DOI:** 10.1101/2025.10.03.680307

**Authors:** Thomas Balan, Mélanie Bazin-Gélis, Marc Guérineau, Valerio Vitali, Coralie Zangarelli, Olivier Arnaiz, Louise Abbou, Abdulwahab Altair, Aménaïde Boutte du Jonchay, Aurélie Camprodon, Marina Giovannetti, Camille Poitrenaud, Emma Schumacher, Julien Bischerour, Anne-Marie Tassin, Vinciane Régnier, Guillaume Chevreux, Sandra Duharcourt, Mireille Bétermier

**Affiliations:** Université Paris Cité, CNRS, Institut Jacques Monod, 75013 Paris, France; Université Paris-Saclay, CEA, CNRS, Institute for Integrative Biology of the Cell (I2BC), 91198, Gif-sur-Yvette, France; SupBiotech Engineering School, 94800 Villejuif, France; Smith College, Bachelor Program in Computer Science and Biological Science, Northampton, Massachusetts, USA; Université Paris Cité, UFR Sciences du vivant, 75205 Paris Cedex 13, France

**Keywords:** SMC, ciliate, transposable element, condensin, endonuclease, programmed DNA elimination

## Abstract

Prokaryotes and eukaryotes use diverse strategies to cope with invading mobile genetic elements, including programmed DNA elimination (PDE). In the ciliate *Paramecium*, elimination of transposable elements and their relics requires the PiggyMac (Pgm) endonuclease and its five PgmL partners, yet how this machinery is targeted to cleavage sites remains unclear. Here, we identified condensin I subunits in the proximity proteomes of Pgm and PgmL4. We show that they belong to a condensin complex that is essential for PDE and localizes to developing somatic nuclei. Depleting the development-specific subunits of this complex blocks DNA elimination, phenocopying a Pgm depletion. Developmental condensin is required for the correct nuclear localization of Pgm and some of the PgmLs. Moreover, Pgm and these PgmLs co-immunoprecipitate with condensin I. Our findings uncover functional and physical interactions between a eukaryotic DNA cleavage machinery and a specialized condensin complex that is critical for PDE in a non-dividing nucleus.

## INTRODUCTION

Mobile genetic elements, such as transposable elements (TEs), viruses or plasmids, exert a profound influence on genome architecture and function^1,2^. They can control gene expression patterns and may compromise genome integrity or stability. They can also benefit their host by promoting the emergence of new genes or cellular functions. Living organisms have evolved various strategies to cope with invading mobile genetic elements^3^. One of the most radical is the elimination of “non-self” DNA, which often involves cleavage by an endonuclease. In bacteria and archaea, diverse machineries can cleave foreign DNA upon entry, such as sequence-based restriction-modification enzymes^4^, RNA-directed CRISPR/Cas nucleases^5,6^ or topology-guided bacterial Wadjet complexes^7,8^. Once integrated, mobile genetic elements can still be removed from their host genomes. Indeed, in some cases, as shown during sporulation in *Firmicutes* bacteria, site-specific excision of a cryptic prophage restores a disrupted regulatory gene^9,10^. During vertebrate lymphocyte differentiation, excision of TE-derived intervening sequences enables the assembly of immunoglobulin or receptor genes^11,12^. In contrast, in an increasing number of multicellular eukaryotes, massive and reproducible programmed DNA elimination (PDE) occurs genome-wide during somatic development, removing variable fractions of germline DNA - in some cases up to 95% - containing TEs, DNA repeats and a few germline-specific genes^13–16^. During PDE, how germline sequences are recognized and targeted for elimination is unclear. The underlying molecular mechanisms likely differ between species and remain largely unknown^17,18^.

Ciliates are valuable microbial models to study PDE during germline/soma differentiation. These unicellular eukaryotes present a nuclear dimorphism, which separates germline and somatic functions between two co-existing distinct nuclei^19^. The diploid micronucleus (MIC) harbors the non-expressed germline genome, which contains multiple TE families^20–24^. During sexual reproduction – conjugation or, in some species, a self-fertilization process called autogamy - the MIC undergoes meiosis and transmits the germline genome to the zygotic nucleus, from which the new germline and somatic nuclei of the next generation are formed (Figure S1). The polyploid somatic macronucleus (MAC) ensures all gene expression. Yet, it is destroyed at each sexual cycle and progeny survival requires that a functional new MAC develops from a copy of the zygotic nucleus. New MAC development involves multiple rounds of genome endoduplication, as well as extensive PDE that removes ∼30% to ∼90% of germline DNA from the somatic genome, depending on species. In *Paramecium tetraurelia*, two types of eliminated sequences have been identified^25,26^. Multi-copy germline TEs, mainly non-LTR retrotransposons and *Tc1/mariner* DNA transposons, are eliminated imprecisely, together with their encompassing regions^20,22^. In addition, ∼45,000 internal eliminated sequences (IESs), originating from ancient *Tc1/mariner* TEs and mostly very short (between 26 and 150 bp), are disseminated throughout the germline genome, often interrupting genes^20,22,24^. Thus, their precise excision is indispensable to assemble a functional version of the somatic genome. PDE involves DNA double-strand breaks (DSBs) at IES ends^27^ and presumably also at the boundaries of imprecisely eliminated sequences^28^. DSBs are introduced by an endonuclease machinery composed of domesticated ancestral PiggyBac transposases - the PiggyMac (Pgm) catalytic subunit and five associated related partners, PgmL1 to PgmL5^29–32^. The broken ends of flanking MAC-destined DNA are then joined by non-homologous end joining (NHEJ) factors^29,33–36^. Coupling between DNA cleavage and DSB repair, mediated by an interaction between Pgm and NHEJ factors, minimizes the risk of errors during genome assembly^37,38^.

Understanding how germline sequences are determined for elimination has been the subject of longstanding research in *Paramecium*. In contrast to classical transposition^39,40^, no strongly conserved sequence that could serve as a Pgm recognition motif is found at IES ends^20,41^. The current view is that small non-coding RNAs corresponding to MIC-restricted sequences absent from the parental MAC^42–45^, drive Polycomb Repressive Complex 2 (PRC2) to their homologous sequences by pairing with nascent transcripts in the developing new MAC^28^. PRC2 marks these sequences by depositing repressive histone marks H3K9me3 and H3K27me3^46–49^. Pgm is thought to be attracted to these marked regions and catalyze their elimination. Available data on the epigenetic control of TE elimination and IES excision are consistent with this model, except for ∼30% of IESs that depend neither on small RNAs nor on histone marks to be excised^22,24,44,47,50,51^. The model does not explain how Pgm is precisely positioned on the boundaries of eliminated DNA and how it is activated to catalyze DNA cleavage.

We have addressed this question by identifying novel factors that associate with the Pgm endonuclease during PDE. We report the discovery of a developmental condensin complex that interacts physically and functionally with Pgm and some of its PgmL partners. Condensins belong to the family of Structural Maintenance of Chromosomes (SMC) complexes that contribute to the spatial organization of DNA and chromatin in all living organisms^52,53^. These multi-subunit complexes drive the intramolecular compaction of chromosome arms through DNA or chromatin loop extrusion. During mitosis, they participate in the individualization of newly replicated sister chromatids and their correct segregation during cell division. Condensins also have non-mitotic functions in interphase nuclei, where they are involved in chromosome organization, the formation of individual chromosome territories, the dispersion of chromocenters or the control of dosage compensation^54^. We show here that the *Paramecium* developmental condensin plays a critical role by assisting Pgm in carrying out PDE and discuss its non-canonical functions in the non-dividing developing new MAC.

## RESULTS

### The Pgm and PgmL4 proximity proteomes include two condensin I subunits

To search for new putative interactants of the Pgm endonuclease, we set up the TurboID *in vivo* proximity labeling technique^55^ in autogamous *Paramecium tetraurelia* cells (Figure S1). We extended our study to the proximity proteome of PgmL4, the phylogenetically closest relative of Pgm, which was shown to interact with Pgm and is also required for DNA elimination^30^. Pgm or PgmL4a fused to the promiscuous biotin ligase TurboID (Pgm-TbID or PgmL4a-TbID, respectively) were each expressed from RNAi-resistant transgenes following microinjection into individual cells (Figure S2A). For each TbID fusion, independent transformants and non-injected controls were allowed to undergo autogamy upon endogenous *PGM* (for Pgm-TbID) or *PGML4a+b* (for PgmL4a-TbID) knockdown (KD). Five hours after the onset of autogamy (T5 time-point), when IES elimination takes place^27,51^ and expression of each fusion transgene is expected to be maximal^56^, *in vivo* protein biotinylation was induced for 1 hour following biotin addition^57,58^. Using fluorescent streptavidin, we observed that biotinylated proteins concentrate in the developing new MACs, coinciding with the localization of Pgm and PgmL4a (Figure S2B). A streptavidin pulldown was performed on whole-cell lysates and among the proteins captured on streptavidin beads, we recovered each TbID fusion, as expected, along with additional proteins that were not detectable in the control samples on western blots (Figures S2C-D). Using specific antibodies, we identified the close paralogs PgmL4a/4b (=PgmL4) among the proteins biotinylated by Pgm-TbID, and Pgm among those biotinylated by PgmL4a-TbID. Following mass spectrometry (MS), quantitative statistical analysis revealed that 94 and 160 protein groups are significantly enriched relative to control samples in the Pgm-TbID and PgmL4a-TbID proximity proteomes, respectively (Figure 1A), with 84 protein groups shared by both datasets (Figure S3, see Zenodo).

**Figure 1.**
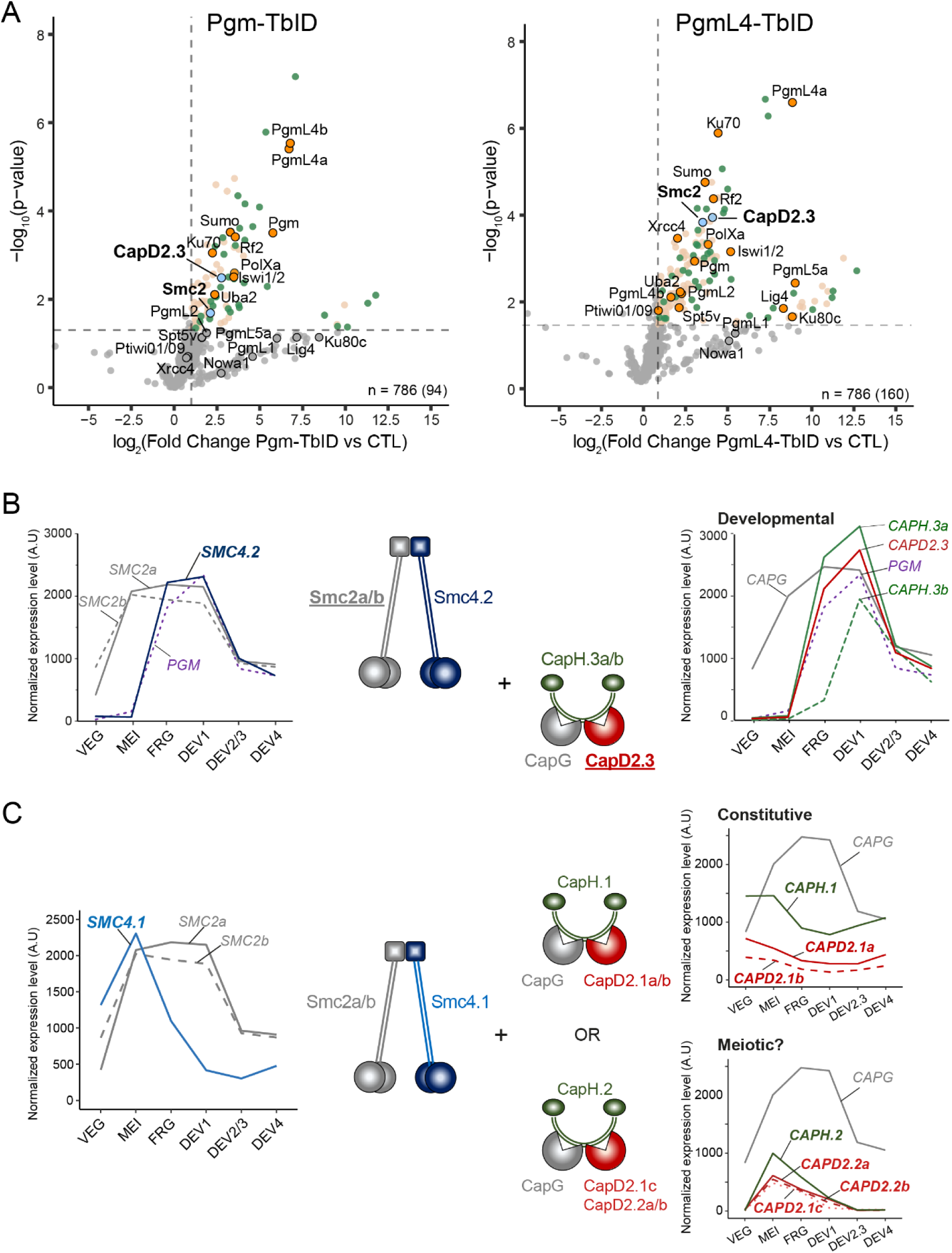
Identification of condensin I subunits in the TurboID (TbID) proximity proteomes of Pgm and PgmL4. (A) Volcano plots of the quantitative mass spectrometry (MS) analysis of protein enrichment in Pgm-TbID (left, 3 replicates) and PgmL4-TbID (right, 4 replicates) experiments. Only protein groups supported by at least 2 unique peptides are shown. Significantly enriched protein groups in the TbID samples over 7 control replicates (p-value ≤0.05 (two-tailed Student t-test), Fold Change (FC) ≥ 2) are represented as colored dots. The total number of protein groups that were identified in each experiment is indicated at the bottom right, with the number of significantly enriched groups in parenthesis. Blue dots: groups containing condensin I subunits; orange dots: groups including proteins involved in PDE; green dots: groups with at least one member expressed from the intermediate or early expression peak cluster. **(B)** Composition of the *P. tetraurelia* developmental condensin I complex. The development-specific subunits (Smc4.2, CapD2.3 and CapH.3a/b) are colored. The subunits identified in the TbID screen (A) are underlined. RNA sequencing profiles of the genes encoding Smc (left) and non-Smc (right) subunits are shown along with *PGM* expression profile (purple dotted curve). **(C)** Inferred composition of the constitutive and putative meiotic condensin I complexes. The Smc2a/b - Smc4.1 heterodimer (left) is proposed to associate with either of two distinct sets of non-Smc regulatory subunits (right) to form a constitutive (top right) or meiosis-specific (bottom right) condensin complex. Expression patterns of the genes encoding each subunit are shown next to each scheme.

As reported previously^56^, the progression of autogamy is accompanied by successive peaks of gene expression. The two proximity proteomes are enriched in proteins encoded by genes from the “early” and “intermediate” expression peak clusters, which include known PDE genes that are induced before or by the time DNA elimination takes place^56,59^ (Figure S3). In particular, and validating our strategy to identify new PDE factors, we found the DNA repair proteins Ku70^35,38^ and PolXa^36^, the RF2 cofactor of the PRC2 complex^48,49^, the chromatin-remodeling factor ISWI1^60,61^ and the Uba2 and SUMO proteins^62^, all of which are essential for PDE. We also identified the Pgm-associated proteins PgmL2 and PgmL5a^30^ and the Lig4/Xrcc4 DNA repair complex^34^ in the PgmL4a-TbID proximity proteome. Among the proteins enriched in both proximity proteomes (Figures 1A and S3), we found homologs of two subunits of condensin I, Smc2 and CapD2^53^.

Eukaryotic condensin I is made of five subunits, the Smc subunits Smc2 and Smc4, the HEAT proteins CapD2 and CapG, and the kleisin CapH, which are encoded by a total of 15 genes in *P. tetraurelia*^63^ (Table S1A). This remarkable expansion results from the combination of segmental and whole genome duplications (WGDs)^64^. Two *SMC2* genes, *SMC2.1* and *SMC2.2*, originating from the most recent WGD^64^ were previously annotated in the *P. tetraurelia* genome^65^. We renamed them *SMC2b* and *SMC2a*, respectively, according to the *Paramecium* gene nomenclature guidelines^66,67^. The corresponding two proteins are 97.54% identical and could not be distinguished in our MS analysis: they will be referred to collectively as Smc2. In contrast, the CapD2 homolog identified in our TbID experiments is clearly encoded by *CAPD2.3*, one of the six *P. tetraurelia CAPD2* genes (*CAPD2.1a/b/c*, *CAPD2.2a*/*b*, *CAPD2.3*) (Table S1A). *P. tetraurelia* also harbors two *SMC4* (*SMC4.1* and *SMC4.2*, which are not WGD paralogs), a single *CAPG* and four *CAPH* genes (*CAPH.1*, *CAPH.2*, *CAPH.3a*/*b*). This multiplicity of genes suggests that there might have been a diversification of condensin I subunits in *P. tetraurelia*, only some of which are found in the proximity proteomes of Pgm and PgmL4a.

### Identification of a development-specific condensin I complex in *P. tetraurelia*

Using published transcriptome data^56^, we examined the expression profiles of the genes encoding all condensin I subunits (Figures 1B-C). *SMC2a/b* and *CAPG* are expressed throughout the life cycle: in vegetative cells, during MIC meiosis and during new MAC development. Thus, we propose that all putative condensin I complexes in *P. tetraurelia* would contain Smc2 and CapG but differ in the other associated subunits. Unique among *CAPD2* genes, *CAPD2.3* belongs to the “intermediate peak” cluster, with its mRNA accumulating specifically during MAC development^56^ (Figure 1B), and is co-expressed with *PGM* and *PGML4a/b*. A similar expression pattern is observed for *CAPH.3a/b* and *SMC4.2*, the latter recently reported to be required for PDE^65^. We therefore speculate that CapD2.3, CapH.3a/b and Smc4.2 assemble with Smc2 and CapG in a developmental condensin I complex, while Smc4.1 and other subunits, possibly CapH1 and CapD2.1a/b, would define a “constitutive” complex present during vegetative growth and throughout cell life cycle (Figure 1C). Consistent with the latter hypothesis, we found that *SMC2* or *CAPG* KDs targeting the constitutive subunit genes induce strong effects during vegetative growth: cells stop dividing and eventually die within a few days (Figure S4A). Similar vegetative phenotypes were documented upon knocking down *SMC4.1*, which is expressed throughout the life cycle of *P. tetraurelia*, with gross defects in MAC segregation during cell division and rapid MIC loss^65^ (Figure S4B-C). Indicative of a ubiquitous role of Smc4.1, a functional Smc4.1-FLAG fusion is detected in MICs and MAC during vegetative growth and in all nuclei during autogamy (Figure S4D-E).

To confirm the composition of the developmental condensin I complex, we sought to identify the protein partners of CapD2.3. Nuclear extracts were prepared at T10 during autogamy from *Paramecium* control cells and from cells expressing a functional tagged CapD2.3-FLAG-HA (Supplementary File S1). First, we conducted nuclear extractions with increasing saline concentrations to determine the conditions where most CapD2.3 is recovered in soluble nuclear fractions (Figure 2A). Western blot analysis indicated that CapD2.3 is exclusively detected in the nuclear fraction, as opposed to the cytoplasmic fraction, consistent with its localization in the new MAC by immunofluorescence (Figure 2A-B). The protein is mostly in the soluble nuclear fraction at 15 mM and 150 mM NaCl (Figure 2A). This extraction profile mirrors that of histone H3, suggesting that CapD2.3 is a chromatin-bound protein (Figure 2A). To identify tight protein interactors of CapD2.3, we performed tandem affinity purification of the FLAG and HA tags from 150 mM NaCl nuclear extracts, followed by quantitative MS, for two replicates in each condition (Figure 2B-C, S5A). Statistical analyses revealed 25 significantly enriched proteins in CapD2.3 tandem immunoprecipitations (IPs) compared with controls (Figure 2C), out of 836 identified proteins. Among the enriched proteins, we recovered CapD2.3, as expected, together with Smc2a, Smc2b, Smc4.2, CapG and CapH.3a. No other condensin subunits were significantly enriched, indicating that CapD2.3 is part of a single condensin I complex. CapD2.3-associated proteins also include proteins involved in PDE: the putative RNA helicase Ema1a^42,48,68^, the chromatin remodeler Iswi1^60,61^, and the DNA-dependent protein kinase DNA-PKcs involved in double-strand break repair^26,69^.

**Figure 2:**
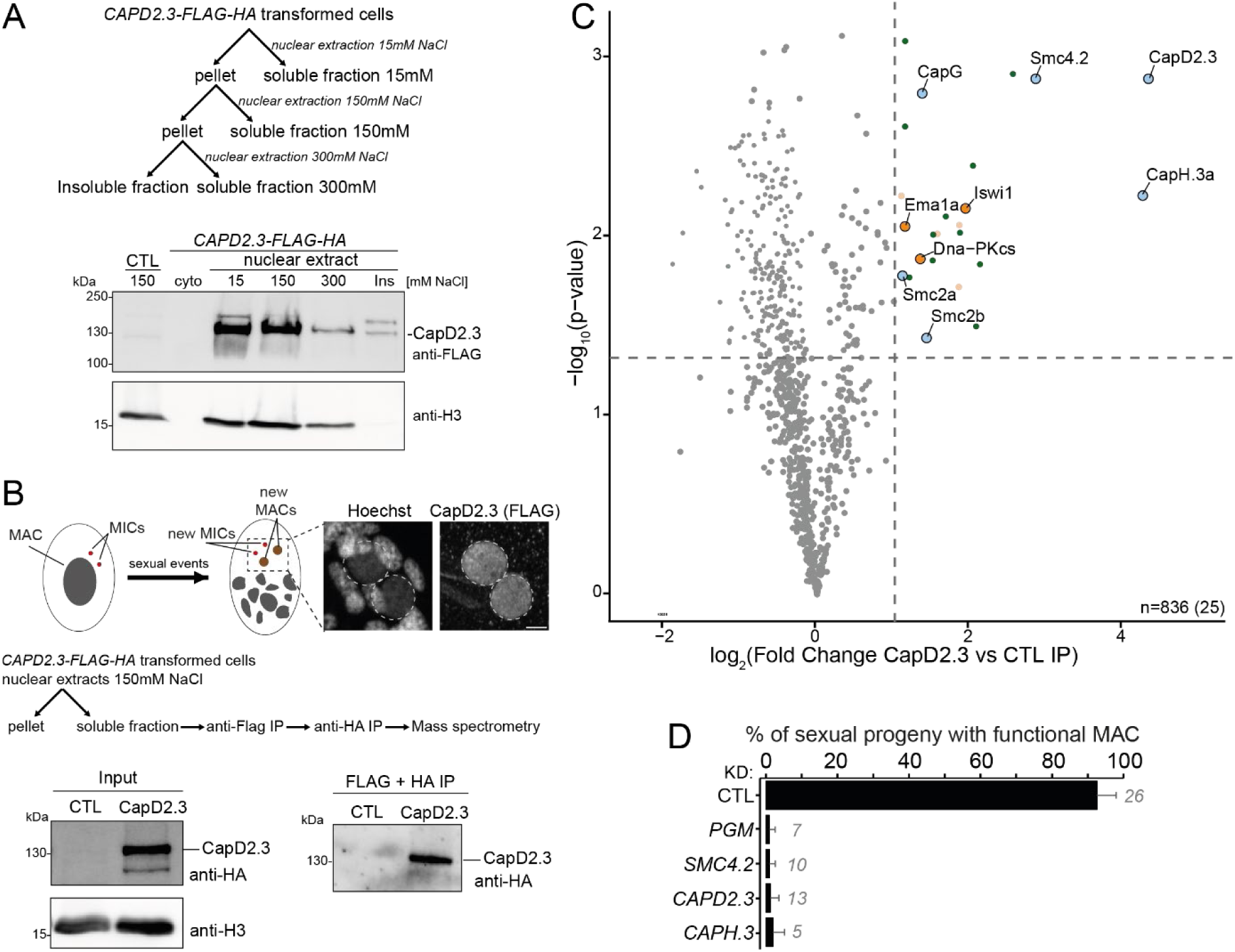
Tandem immunoprecipitation of CapD2.3 identifies the developmental condensin I complex. (A) (Top) Schematic representation of serial nuclear extracts at increasing salt concentrations. (Bottom) Western blot analysis of CapD2.3-FLAG-HA protein levels from nuclear extracts prepared at increasing salt concentrations from cells expressing a *CAPD2.3-FLAG-HA* transgene and from control non-injected cells (CTL), at T10 during autogamy. Bottom lane shows histone H3 levels. **(B)** (Top) Schematic representation of CapD2.3 tandem immunoprecipitation experiments, with anti-FLAG immunostaining of cells transformed with a *CAPD2.3-FLAG-HA* transgene, at T10 during autogamy (top right panels). Developing new MACs are circled with a dotted line. Scale bar: 5 μm. (Bottom) Western blot analysis of nuclear extracts from wild-type cells (CTL) or cells expressing a CapD2.3-FLAG-HA functional protein (CapD2.3) before (input) or after tandem affinity purification (FLAG +HA IP). Anti-HA antibodies are used to detect CapD2.3 and anti-histone H3 antibodies for normalization. Predicted MW for CapD2.3-FLAG-HA: 138.2 kDa. **(C)** Volcano plot of the quantitative label-free mass-spectrometry analysis of CapD2.3-FLAG-HA tandem affinity purification. Significantly enriched proteins in CapD2.3 tandem IP (2 replicates) over control IP (2 replicates). p-value ≤ 0.05 (two-tailed Student t-test), Fold Change (FC) ≥ 2, only proteins identified with more than 5 unique peptides are considered. The total number of identified proteins is indicated at the bottom right, with the number of significantly enriched proteins in parenthesis. Components of the developmental condensin I complex are highlighted in light blue. Proteins known to be involved in PDE are highlighted in orange. Proteins expressed from the intermediate or early expression peak clusters are highlighted in dark green. **(D)** Survival of post-autogamous progeny with a functional new MAC following *ND7* (CTL), *PGM*, *SMC4.2*, *CAPD2.3* or *CAPH.3* KD. The number of independent experiments is indicated on the right of each bar. Error bars correspond to the standard deviation (SD) of the mean % of cells in each condition.

*In vivo* interaction of CapD2.3 and Smc4.2 was confirmed by co-IP using soluble nuclear protein extracts prepared at T5-T10 from autogamous cells co-expressing CapD2.3-HA and Smc4.2-FLAG (Figure S5B, see SupInfo file). Endogenous *CAPD2.3* and *SMC4.2* genes were knocked down in these experiments, since depleting each endogenous protein increases the nuclear amounts of its tagged counterpart (Figure S5C). Indicative of an interaction between the two tagged proteins, we observed that CapD2.3-HA co-precipitated with Smc4.2-FLAG in the presence of anti-FLAG antibodies (Figure S5D). Reciprocally, Smc4.2-FLAG co-precipitated with CapD2.3-HA in the presence of anti-HA antibodies, albeit with lower efficiency. We conclude that the *Paramecium* developmental condensin I complex is composed of the Smc proteins Smc2a/b and Smc4.2 and of the non-Smc subunits CapG, CapH.3a and CapD2.3.

### The development-specific condensin I is essential for MAC development

The presence of subunits of the developmental condensin I complex in the proximity proteomes of Pgm and PgmL4a prompted us to carry out functional assays using gene knockdowns (KDs). We focused on the genes encoding the three development-specific subunits CapD2.3, Smc4.2 and CapG, because the strong vegetative phenotypes observed upon depleting Smc2 or CapG (Figure S4) make it difficult to investigate their role during autogamy. Knocking down any of the three development-specific condensin genes (*CAPD2.3, SMC4.2* or *CAPH.3*) resulted in the absence of viable post-autogamous sexual progeny, similar to a *PGM* KD, indicating that the new MACs that develop under these conditions are not functional (Figure 2D). To examine the localization of the three developmental subunits, we expressed tagged CapD2.3-HA, Smc4.2-FLAG and CapH.3a-FLAG fusions from functional RNAi-resistant transgenes in cells knocked down for each respective endogenous gene, and performed immunostaining of cells, during vegetative growth and at T5-T10 during autogamy. Similar to Pgm, and in agreement with a previous report on Smc4.2^65^, we detected each tagged protein excusively in the developing new MACs of autogamous cells (Figure 3A-C and S6A). Their specific nuclear localization is consistent with the three subunits belonging to the same condensin I complex involved in new MAC development (Figure 2C-D). In addition, we observed a significant decrease in the nuclear immunofluorescence signal of each tagged protein upon depletion of any of the other subunits (Figure 3D-F and S6B), while no effect on the steady-state levels of each fusion protein in whole cell extracts was detected by western blot analysis (Figure 3D-F). This indicates that the three development-specific condensin I subunits strongly depend on each other for their stable localization in the new MACs. In contrast, we found that they localize normally in the new MAC upon *PGM* KD (Figure 3D-F).

**Figure 3:**
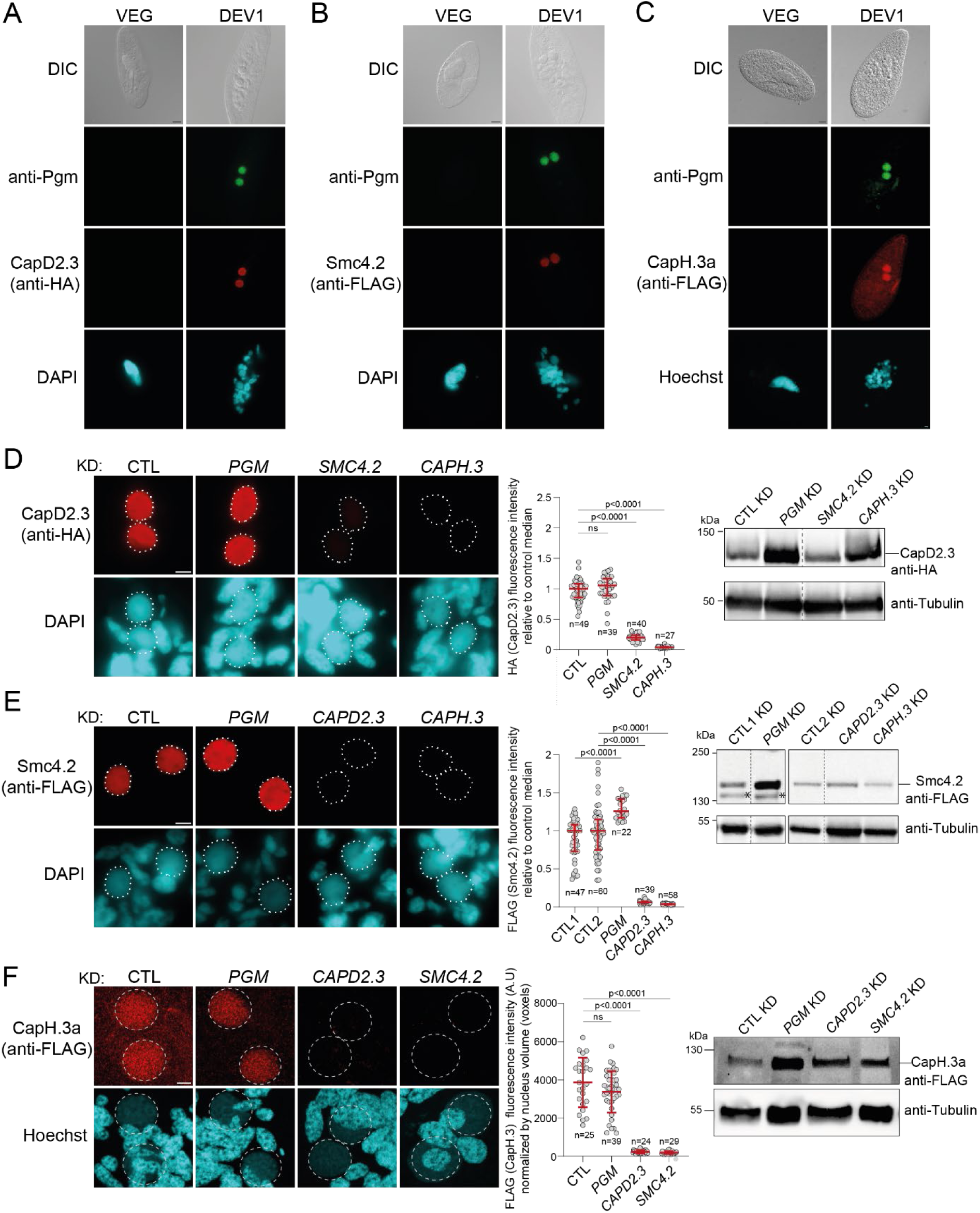
Subcellular localization and *in vivo* inter-dependence of developmental condensin I subunits. (A-C) Immunostaining of Pgm and CapD2.3-HA (A), Smc4.2-FLAG (B) or CapH.3a-FLAG (C) in vegetative (VEG) and autogamous cells collected at T5-T10 (DEV1, see Figure S1), visualized using epifluorescence (A,B) or confocal microscopy (C). New developing MACs are labeled with anti-Pgm antibodies. Panel A corresponds to cells subjected to control *ND7* KD and a qualitatively similar localization of CapD2.3-HA is observed in a *CAPD2.3* KD (Figure S6A). Panels B and C show cells upon *SMC4.2* and *CAPH.3* KDs, respectively. Differential interference contrast (DIC) images are displayed on top. Scale bar: 10 µm. **(D-F)** Nuclear localization of CapD2.3-HA (D), Smc4.2-FLAG (E) or CapH.3a-FLAG (F) in autogamous cells depleted of Pgm or development-specific condensin subunits (T5-T10). (Left) Immunostaining of CapD2.3-HA, Smc4.2-FLAG or CapH.3a-FLAG in nuclei upon *ND7* (D,E) *or ICL7* (F) control KDs (CTL), *PGM*, *SMC4.2*, *CAPD2.3* or *CAPH.3* KDs, visualized using epifluorescence (D,E) or confocal (F) microscopy. Developing new MACs are circled with a dotted line. Scale bar: 5 µm. (Middle) HA (CapD2.3) or FLAG (Smc4.2) fluorescence intensities in developing new MACs in each KD, divided by the median fluorescence intensity in their respective control KD (D,E), and FLAG (CapH.3a) fluorescence intensities in developing new MACs normalized to nucleus volume (Figure S6B) (F). The number of nuclei is indicated for each sample. Bars correspond to the median and 1^st^ and 3^rd^ quartiles of each distribution (D,E) or to the mean ± SD (F). *p*-value thresholds are indicated (not significant (ns): p> 0.05. Mann-Whitney U statistical tests, with Bonferroni correction for panels D-E). (Right) Western blot analysis of CapD2.3-HA, Smc4.2-FLAG or CapH.3a-FLAG in whole-cell extracts from DEV1 cells under the same conditions. Asterisks in panel E indicate a non-specific band that is occasionally detected with anti-FLAG antibodies, even in the absence of the transgene (full membrane in Zenodo).

### The developmental condensin I complex is required for programmed DNA elimination

Given the specific localization of developmental condensin I subunits in the new MACs and the absence of viable sexual progeny caused by *SMC4.2*, *CAPD2.3* or *CAPH.3* KDs, we investigated whether PDE is properly completed under these conditions by deep-sequencing the DNA extracted from new MACs at a late developmental stage. We selectively sorted the developing new MACs from cells subjected to *SMC4.2*, *CAPD2.3*, *CAPH.3a+b*, *PGM* or control KDs at T28-T30 during autogamy, when IES excision has been completed and imprecise TE elimination is ongoing^51^ (Figure S7A-C). Following DNA purification and deep sequencing, we found that *SMC4.2*, *CAPD2.3* or *CAPH.3a+b* KDs induce the full retention of all 44,928 known IESs in the new MAC, just like a *PGM* KD (Figure 4A). Compared to the controls, where TE elimination is already well advanced – yet not complete - at this stage, depleting any of the three development-specific subunits of condensin I also affects TE elimination, similarly to depleting Pgm (Figure 4B). Thus, depletion of the development-specific subunits of condensin I results in the same inhibitory effect on PDE as depleting the Pgm endonuclease itself.

**Figure 4.**
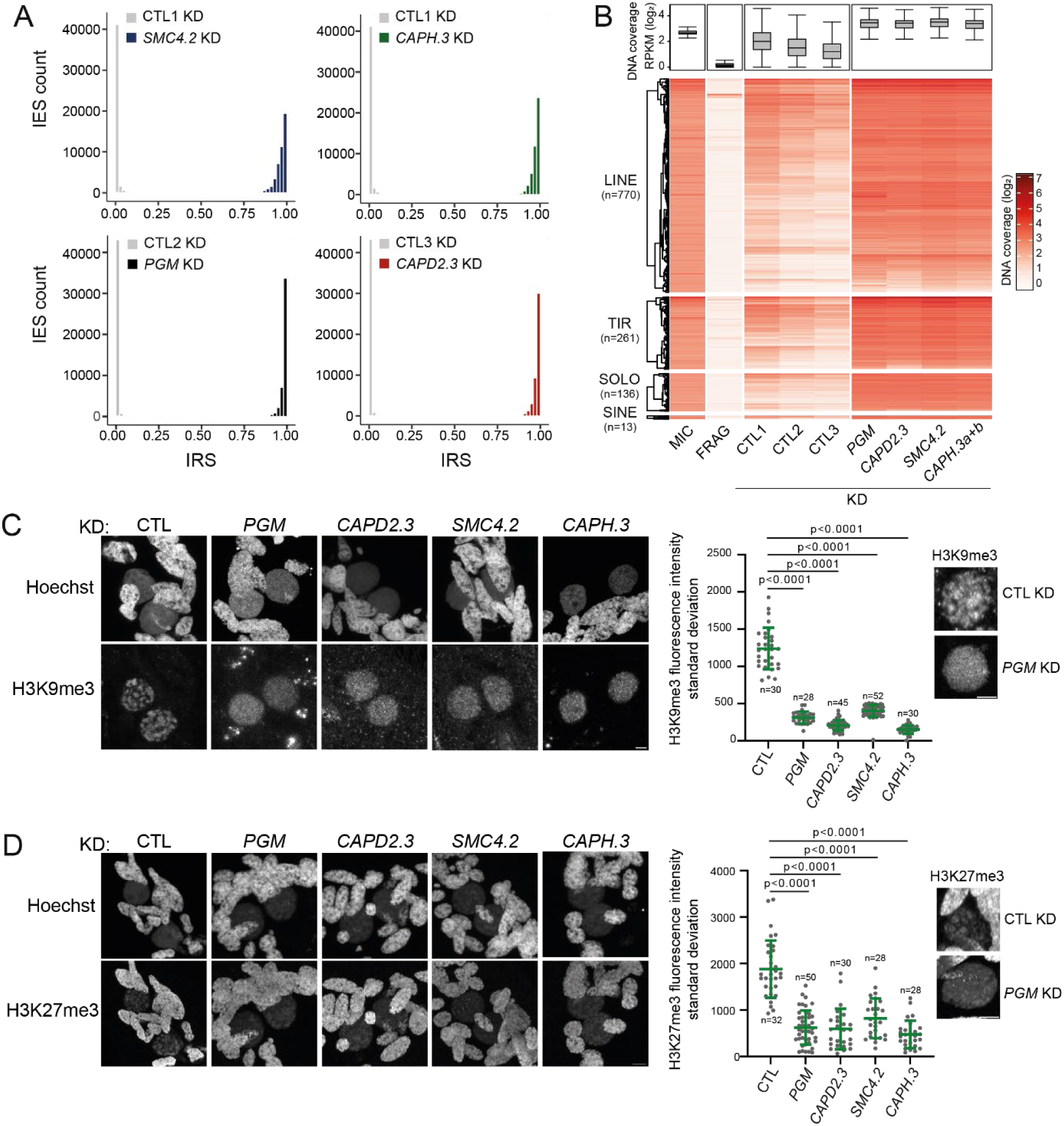
Developmental condensin I is required for genome-wide DNA elimination. (A) Distribution of IES retention scores in sorted developing new MACs collected at T28-T30 from autogamous cells subjected to *SMC4.2, CAPH.3a+b, PGM* or *CAPD2.3* KD. Histograms for each corresponding control (CTL1-3: Empty vector) are shown in gray. A bin width of 0.02 was used to calculate IES frequencies. **(B)** Heatmap of normalized TE DNA coverage in the same samples as in A. TE copies are grouped according to their families. The coverage distribution (RPKM: Reads Per Kilobase per Million mapped reads) for all TE copies is shown as a boxplot on top (log2 scale). FRAG: DNA from purified MAC fragments^51^. MIC: DNA from flow-cytometry-sorted micronuclei^22^. LINEs: class I Long Interspersed Nuclear Elements. TIRs: class II DNA transposons with Terminal Inverted Repeats. SINEs: class I non-autonomous Short Interspersed Nuclear Elements. SOLO: class I non-autonomous elements carrying ORF1 only. **(C-D)** Immunostaining of H3K9me3 (C) and H3K27me3 (D) in wild-type cells at T10 upon *ICL7* (CTL), *PGM*, *CAPD2.3*, *SMC4.2* or *CAPH.3* KD visualized using confocal microscopy. Developing new MACs are circled with a dotted line. Scale bar: 5 μm. Quantifications of the standard deviation of H3K9me3 (C) and H3K27me3 (D) fluorescence intensity in the new MACs are shown for each KD. The number of nuclei is indicated. Bars correspond to mean ± SD. Mann-Whitney statistical tests. The inset displays representative H3K9me3 and H3K27me3 immunostaining images upon *ICL7* (CTL) and *PGM* KD in the new MACs. Scale bar: 5 μm.

We previously showed that depletion of Pgm affects the localization pattern of H3K27me3 and H3K9me3 modifications in the new MACs^47^. While H3K27me3 and H3K9me3 normally concentrate into nuclear foci, these foci no longer form in Pgm-depleted cells. To determine the consequences of condensin I depletion on the accumulation and distribution of H3K9me3 and H3K27me3, we performed immunofluorescence with anti-H3K9me3 and anti-H3K27me3 antibodies^46^ during autogamy (T10). In control cells, as expected^46,47^, H3K27me3 is detected in the maternal MAC and in the developing new MACs, while H3K9me3 only accumulates in the developing new MACs. In cells subjected to *CAPD2.3*, *SMC4.2*, or *CAPH.3* KDs, the H3K9me3 and H3K27me3 fluorescence intensities in the developing new MACs are similar to those in the control (Figure S7D-E), yet no nuclear foci are visible, similarly to what is observed in a *PGM* KD (Figure 4C-D). Quantification of the standard deviation of H3K9me3 and H3K27me3 fluorescence intensity confirmed the lack of histone modification foci in the developing new MAC. In conclusion, developmental condensin I, like Pgm, is essential for PDE and for the formation of H3K9me3 and H3K27me3 nuclear foci in the new MACs.

### Developmental condensin I stabilize the Pgm/PgmL machinery within the new MACs

Because depletion of the developmental condensin I appears to phenocopy depletion of Pgm, we used specific custom primary antibodies (Figure S8 and S9) to examine the effect of condensin I depletion on the nuclear localization of Pgm and each of its five PgmL partners^30^. As a control, we first confirmed on western blots that Pgm and PgmLs are present in whole cell extracts during autogamy (T5-T10) upon *CAPD2.3*, *SMC4.2* or *CAPH.3* KD (Figure S10A). We then permeabilized, fixed and immunolabeled autogamous cells at the same time-points (Figure 5A). Quantitative analysis of the fluorescence signal in the developing new MACs revealed a decrease of Pgm and PgmL4 immunofluorescence intensities upon *CAPD2.3*, *SMC4.2* or *CAPH.3* KD compared to control conditions, indicating that the nuclear localization of both proteins is strongly affected. Immunofluorescence intensities also dropped for PgmL2 and PgmL5. In contrast, little or no decrease was observed for PgmL1 and PgmL3, with a more marked effect of *SMC4.2* KD relative to the other two KDs. These observations suggest that the developmental condensin I complex is necessary for PgmL2, PgmL4, PgmL5 and Pgm to correctly localize in the developing new MACs, which may explain the strong effect of *SMC4.2*, *CAPD2.3* and *CAPH.3* KDs on PDE. Notably, the dependence relationship between the Pgm/PgmL proteins and the developmental condensin I complex does not appear to be reciprocal since depletion of Pgm has no effect on the localization of Smc4.2, CapD2.3 and CapH.3a in the developing new MACs (Figure 3D-E).

**Figure 5.**
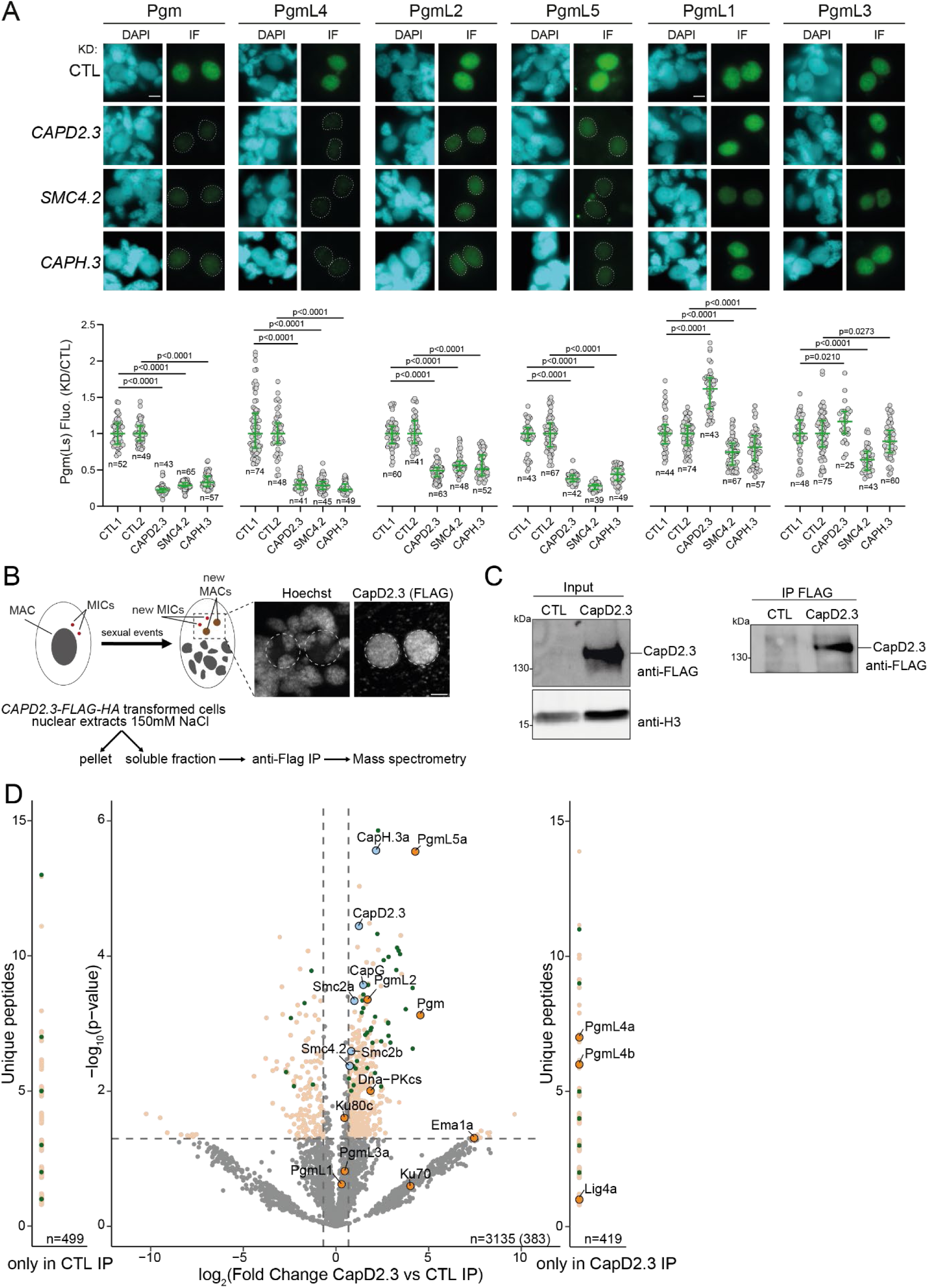
Functional and physical interaction of the developmental condensin I complex with Pgm and its PgmL partners. (A) Pgm and PgmL nuclear localization in cells upon control (CTL: Empty vector), *CAPD2.3*, *SMC4.2* or *CAPH.3* KD. Pgm and PgmLs were immunostained at T5-T10 during autogamy, following pre-extraction to visualize stable proteins using epifluorescence microscopy. (Top) Visual fields restricted to the two developing new MACs. Images are representative of the median fluorescence intensity of the cell population for each condition. Scale bar: 5 µm. (Bottom) Quantification of Pgm or PgmL fluorescence intensities in new developing MACs under each condition, divided by the median fluorescence intensity in their respective CTL KD. The number of imaged nuclei is indicated. Two CTL samples were analyzed: CTL1 for *CAPD2.3* and *SMC4.2* KD, CTL2 for *CAPH.3a+b* KD. Bars correspond to the median and 1^st^ and 3^rd^ quartiles of each distribution. Mann-Whitney U statistical test with Bonferroni correction. n.s: non-significant. **(B)** Schematic representation of CapD2.3 immunoprecipitation experiments. Anti-FLAG immunostaining of a cell expressing a *CAPD2.3-FLAG-HA* transgene, at the time when nuclear extracts were prepared (T10). Developing new MACs are circled with a dotted line. Scale bar: 5 μm. **(C)** Western blot analysis of nuclear extracts from cells expressing a *CAPD2.3-FLAG-HA* transgene and non-injected cells (CTL) before (input) and after affinity purification (IP FLAG). Anti-FLAG antibodies are used to detect CapD2.3 and anti-H3 antibodies for normalization. **(D)** Volcano plot of the quantitative label-free mass spectrometry analysis of CapD2.3-FLAG-HA affinity purification. Significantly enriched proteins in CapD2.3 IP (3 replicates) over control IP (3 replicates), p-value ≤ 0.05 (two-tailed Student t-test), Fold Change (FC) ≥ 2. Proteins only detected in CapD2.3 or in control IP (CTL) are displayed on either side with the number of unique peptides detected. The condensin complex (light blue), the Pgm machinery (orange) and proteins expressed from the intermediate or early expression peak clusters^56^ are highlighted in dark green.

### The developmental condensin I complex physically interacts with Pgm and some of its PgmL partners

To determine whether Pgm or its PgmL partners interact *in vivo* with the developmental condensin I complex, we performed CapD2.3 IPs from soluble nuclear extracts of control cells and cells expressing FLAG-HA-tagged CapD2.3 at T10 in triplicates, followed by quantitative MS (Figure 5B-C, S10B). Here, unlike the tandem affinity purification shown in Figure 2, we used only a FLAG tag to immunoprecipitate CapD2.3. This less stringent approach was chosen to identify proteins associated with the developmental condensin I complex. Statistical analyses revealed 383 significantly enriched proteins in CapD2.3 IP compared with controls, out of 3135 identified proteins (Figure 5D). We recovered all components of the developmental condensin I complex (Smc2a and b, Smc4.2, CapG, CapH.3a and CapD2.3), as expected. We also identified Pgm, PgmL2 and PgmL5a among the enriched proteins. We found PgmL4a and b among the proteins that are only detected in the CapD2.3 IPs, but not in the control IPs. In contrast, PgmL1 and the PgmL3 paralogs were not enriched in CapD2.3 IP, suggesting there is no physical association between these proteins and the condensin I complex (Figure 5D). This is consistent with the lack of effect of condensin I complex depletion on PgmL1 and PgmL3 localization (Figure 5A). In addition to Ema1a^42,48,68^ and Dna-PKcs^26,69^, which were among the CapD2.3-associated proteins in the tandem IP (Figure 2), we also identified the NHEJ component Lig4a^34^ in CapD2.3 IP (Figure 5D). Thus, the condensin I complex physically interacts, either directly or indirectly, with the Pgm endonuclease machinery at the time of DNA elimination.

## DISCUSSION

In this study, we identified two subunits of condensin I, CapD2.3 and Smc2, in the TurboID proximity proteomes of the Pgm endonuclease and of one of its known partners, PgmL4a. Tandem affinity purification of CapD2.3 established that it is a component of a development-specific condensin I complex containing Smc2a/b and three other subunits, Smc4.2, CapH.3a and CapG. This complex localizes in the developing new MAC by the time DNA elimination takes place, and depleting any of its development-specific subunits inhibits PDE, phenocopying a Pgm depletion. We show that the developmental condensin I is required for the localization of Pgm and some of its known PgmL co-factors - PgmL2, 4 and 5 - in the developing new MAC. Consistent with this functional interaction, Pgm and the same PgmL subset immunoprecipitate with the developmental condensin I complex from nuclear extracts. We discuss how the interaction between condensin I and the *Paramecium* DNA elimination machinery may promote productive PDE.

### Diversification of condensin I complexes in *Paramecium*

Four SMC complexes have been described in eukaryotes: cohesin, condensin I, condensin II and the SMC5/SMC6 complex^53^. Ciliates have lost condensin II and the SMC5/SMC6 complex, but they all have cohesin and condensin I^70–72,63^. In *P. tetraurelia* and *T. thermophila*, there has been an expansion of the genes encoding the condensin I subunits CapH and CapD2 (Table S1) ^63,71,72^. In addition, while *T. thermophila* has a unique *SMC4* gene^73^, *P. tetraurelia* has two paralogs^65^. Such gene expansion could have enabled the diversification of condensin I complexes by broadening the possible combinations of subunits with diverse expression patterns and nuclear localization. As suggested for *T. thermophila*^71,72^, different condensin I complexes may fulfill distinct functions in the MIC and the MAC, during vegetative growth or sexual processes.

Based on transcriptome data^56^ and the KD phenotypes that we (this study) and others^65^ have observed, at least three condensin I complexes may exist in *P. tetraurelia*, depending on which subunits associate with Smc2 and CapG (Figure 1B-C). A constitutive condensin I complex, present at all stages of the life cycle and the only active complex during vegetative growth, is composed of Smc2, CapG, Smc4.1, CapH.1 and CapD2.1a/b. It is required for MAC division (Figure S4), as reported in previous studies of Smc4 homologs in *Tetrahymena*^73^ and *P. tetraurelia*^65^. In *Tetrahymena*, condensin I is necessary to maintain the multiple copies of each MAC “chromosome” in separate territories and prevent them from clustering together, which may avoid chromosome loss upon MAC amitotic division^71^. Given the phenotypes observed upon *SMC2*, *SMC4.1* and *CAPG* KDs (Figure S4), a similar function may be proposed for the *Paramecium* constitutive condensin I during amitotic division of the vegetative MAC.

The constitutive condensin I complex is also required for MIC maintenance, since *SMC4.1* KD triggers MIC loss within a few vegetative fissions. MIC loss would prevent the formation of gametic nuclei when surviving cells engage in sexual reproduction. As a consequence, no zygotic nucleus - hence no new MAC - would be formed in the progeny, which might explain previously reported phenotypes following autogamy of Smc4.1-depleted cells, with no detectable IES retention but strong lethality in sexual progeny^65^. During meiosis and/or early stages of the sexual cycle, another specific condensin I complex, composed of Smc2, Smc4.1, CapG, CapH.2, CapD2.1c or CapD2.2a/b, may be active (Figure 1C). Further studies will help to confirm the existence of the constitutive and meiotic condensin I complexes in *Paramecium* and address their roles in the different nuclei.

Here, we have characterized a developmental condensin I complex comprising five components: the constitutive subunits Smc2 and CapG, and the three development-specific subunits Smc4.2, CapD2.3 and CapH.3a, which depend upon each other for their correct localization in the developing new MAC (Figure 3). Interestingly, the genes encoding all subunits of this development-specific condensin I, including *SMC2a/b* and *CAPG*, are the only condensin genes to become overexpressed at late autogamy stages when PDE is impaired upon *PGM*, *KU80c* or *XRCC4* KDs (Figure S7F). This observation lends further support to our previous suggestion that novel PDE genes may be found within the subset of genes from the “early” and “intermediate” expression peaks that are aberrantly upregulated when PDE does not proceed normally^59^. In another study, a co-IP search for Smc4.2 partners only retrieved Smc2, no other condensin I subunit was found^65^. This discrepancy with our data might be explained in part by the harsher lysis conditions used by Zhang *et al* for the preparation of soluble whole-cell extracts, which may have caused the dissociation of the complex. We established that the CapD2.3-associated developmental condensin I complex contains Smc4.2, but not Smc4.1 (Figure 2), which confirms the existence of a development-specific condensin I complex in *Paramecium*.

### The developmental condensin I complex interacts with Pgm and a subset of its PgmL partners

Since CapD2.3 and Smc2 are enriched in the proximity proteome of Pgm-TbID, we infer that the condensin I complex is spatially close to Pgm during PDE. Our co-IP data demonstrate that the entire developmental condensin I complex associates with Pgm during new MAC development (Figure 5). Future work will confirm whether this physical association is mediated by direct protein-protein contacts or established through an intermediate molecule. Moreover, we provide evidence for a functional interaction between Pgm and developmental condensin I, since Pgm requires condensin I to stably localize in the developing new MAC. In contrast, the nuclear localization of condensin I is independent of Pgm (Figure 3). Finally, we show that the KD of individual condensin I subunit genes phenocopies a *PGM* KD and interferes with the formation of a functional new MAC (Figure 4). DNA elimination is severely compromised genome-wide, with 100% retention of all IESs and inhibition of TE elimination (Figure 4), which in turn likely impairs chromosome fragmentation, as discussed previously^25^. In *Tetrahymena* as well, a specialized condensin I complex participates in new MAC development^72^. However, its contribution to PDE seems somewhat different from that of its *P. tetraurelia* counterpart. Indeed, the knockout of *Tetrahymena* condensin I genes blocks IES and TE elimination, while still allowing for chromosome breakage and *de novo* telomere addition. Moreover, co-IP or immunolocalization experiments provided no evidence that the *Tetrahymena* condensin I interacts with Tpb2p, the endonuclease responsible for DNA elimination in this ciliate^72^.

We reported previously that Pgm works with five related PgmL co-factors, which are all necessary for PDE and the correct localization of Pgm in the developing new MAC^30^. Our TurboID data reveal that 89% of the Pgm proximity proteome is included in that of PgmL4a. The significant enrichment of Pgm and PgmL4 in the two proteomes (Figure 1) suggests that these proteins are close to each other during PDE. Moreover, several lines of evidence establish a distinction between two sets of subunits that compose the Pgm/PgmL machinery, based on their interaction with the developmental condensin I complex. Indeed, the depletion of condensin I not only interferes with the nuclear localization of Pgm, but also with that of PgmL2, PgmL4 and PgmL5 (Figure 5A). In contrast, the localization of PgmL1 and PgmL3 in the developing new MAC is barely affected. Consistent with these observations, Pgm, PgmL2, PgmL4 and PgmL5 co-precipitate with the condensin I complex, while PgmL1 and PgmL3 do not (Figure 5B). The distinction between the two sets of Pgm/PgmL subunits is reminiscent of our previous report that depleting either Pgm, PgmL2, PgmL4 or PgmL5 completely abolishes PDE, while low residual IES excision activity is still detected in PgmL1- or PgmL3-depleted cells, with errors in the selection of excision boundaries^30^.

In conclusion, our data indicate that the *Paramecium* developmental condensin I complex is a key player in PDE. We propose that the DNA elimination machinery of *P. tetraurelia* is composed of two distinct sets of subunits, which are all essential: a condensin I-interacting set made of Pgm, PgmL2, PgmL4 and PgmL5 on the one hand, and a condensin I-independent set comprising PgmL1 and PgmL3 on the other (Figure 6A).

**Figure 6.**
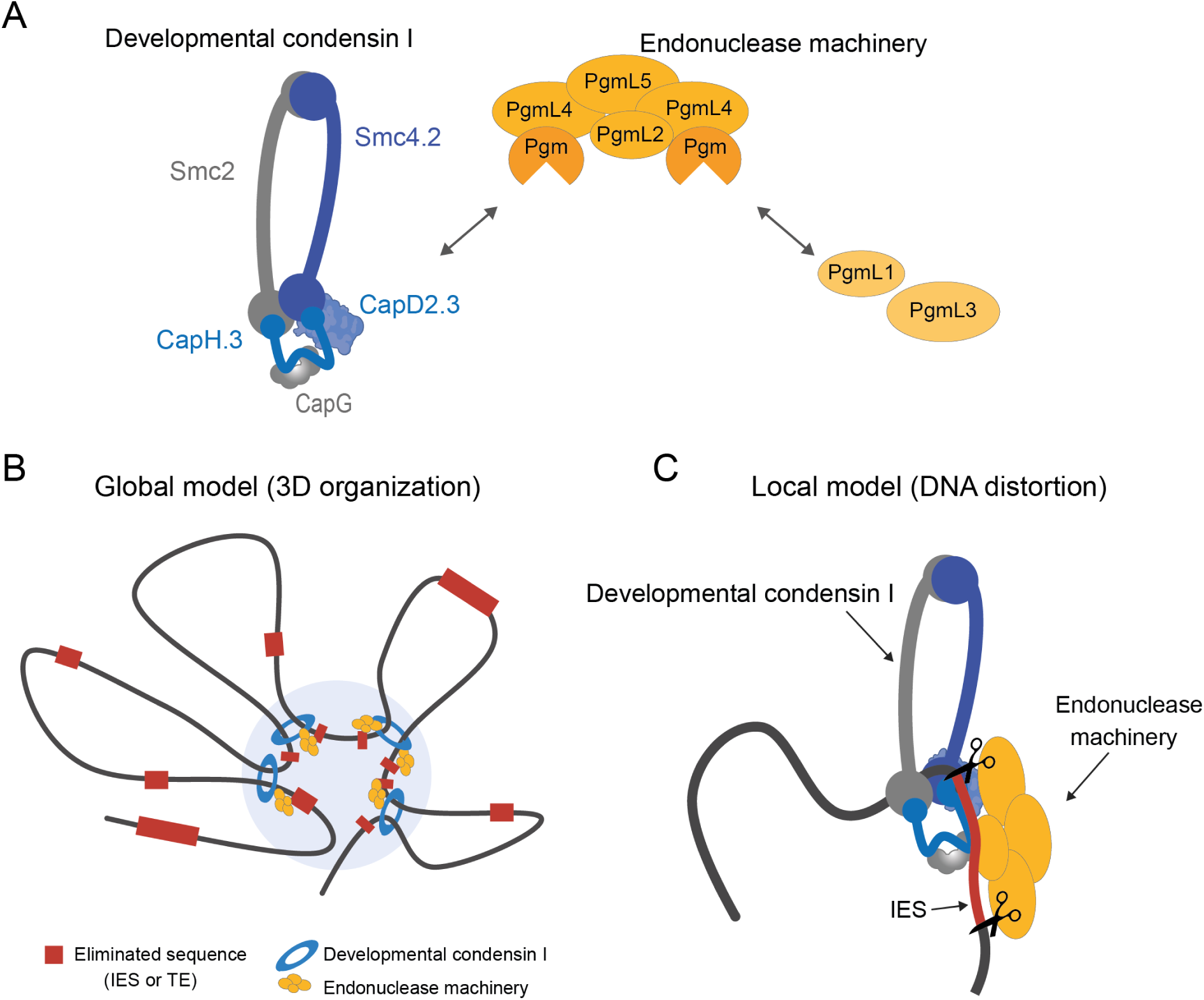
Models of condensin-assisted DNA elimination in *Paramecium*. **(A)** Interaction of developmental condensin I (shown in blue and gray) with components of the Pgm endonuclease machinery (shown in orange). This study reveals that condensin I interacts functionally and physically with Pgm, PgmL2, PgmL4 and PgmL5. It is still unclear whether these four subunits form a complex and, if so, what the organization and stoichiometry of this putative complex would be. PgmL1 and PgmL3 were not found to interact with condensin. However, they are required to stably anchor Pgm in the developing new MAC and allow it to carry out PDE^30^. This suggests that they also interact with the first set of Pgm/PgmLs. Double-headed arrows represent the interactions involving Pgm and its PgmL partners that have been uncovered so far. **(B)** 3D organization of DNA elimination compartments. In this global model, condensin-mediated DNA loop extrusion is proposed to bring together germline DNA sequences that will be eliminated at the same developmental stage by the Pgm/PgmL endonuclease complex. **(C)** Local DNA distortion. Upon DNA binding or during the process of loop extrusion, the condensin complex may locally bend the DNA and favor the correct positioning of the Pgm/PgmL catalytic site on its cleavage sites. The two models are not mutually exclusive.

### A non-canonical role of the developmental condensin I during PDE in *Paramecium*

What could be the role of *Paramecium* condensin I in PDE? In line with other reports^73,71,72,65^, our results in *Paramecium* confirm that ciliate condensin I complexes have a nuclear localization. During the sexual cycle, new MAC development involves a succession of genome endoduplication rounds^51^. However, while their genome undergoes PDE, and until they reach their final ploidy, the developing new MACs do not divide (Figure S1). During endoreplication, developmental condensin I might be involved in the separation of newly replicated sister chromatids and/or the formation of individual chromosome territories, as proposed above for constitutive condensin I. However, we observed no defect in DNA amplification within the developing new MACs of condensin I-depleted cells (Figure S6 and S7). Therefore, the absence of a general developmental arrest suggests that the developmental condensin I has another more specific function during new MAC development.

We show here that condensin I is required upstream of Pgm-mediated DNA cleavage. At this stage, we cannot formally distinguish between a role in the control of Pgm nuclear localization, or in the stabilization/assembly of the Pgm-associated machinery. We may however propose two non-exclusive hypotheses for the role of developmental condensin I that would take into account its interaction with the DNA elimination machinery. In a first “global” model, condensin-mediated loop extrusion may drive the compartmentalization of MIC-limited versus MAC-destined sequences in the new developing MAC before the start of PDE (Figure 6B), a phenomenon that has recently been reported in other ciliates^74,75^. This would resemble the role of cohesin in the formation of intrachromosomal topologically-associating domains^76,77^ or the contraction of genomic loci involved in the recombination of immunoglobulin genes^78,79^. By interacting with Pgm and its partners, condensin I may recruit the DNA elimination machinery to the compartmentalized MIC-limited regions. Consistent with this model, diverse and dynamic nuclear foci involving eliminated chromatin structures have been described during new MAC development in *P. tetraurelia*, such as those formed by H3K9me3/H3K27me3-marked chromatin^46,47^ (Figure 4B) or CenH3-associated centromeric chromatin^80^. There is, however, no strict colocalization of these chromatin foci with Pgm or condensin I, and a detailed understanding of the interplay between spatial genome organization and DNA elimination will require further investigation. A second “local” model may explain the essential role of condensin I in the excision of all 45,000 IESs (Figure 6B). Indeed, previous observations suggested that both ends of an IES establish a crosstalk before excision, which may involve torsional constraints on DNA^20,81^. In addition, we show here that the chromatin remodeler Iswi1/2, identified in the proximity proteomes of Pgm and PgmL4, copurifies with the developmental condensin I complex (Figure 2C). Application of our quantitative analysis pipeline to Iswi1-co-IP data published by others^61^ confirms that Iswi1 co-immunoprecipitates with several condensin I subunits and a subset of PgmL proteins (see Zenodo). Because Iswi1 is thought to decrease the density of nucleosomes on IES DNA, this interaction suggests that condensin I may position the PDE machinery on nucleosome-poor DNA. During the process of loop extrusion, condensins were shown to induce sharp distortions and topological changes on DNA^82,83^. We propose that, in *Paramecium*, condensin I-induced local changes in DNA conformation may favor the precise positioning of the associated Pgm endonuclease on its cleavage sites at IES boundaries. This would be reminiscent of the bacterial Wadjet system, where an SMC-related complex activates DNA cleavage by an associated endonuclease subunit, by bending or twisting the DNA molecule during loop extrusion^84,85^. Our finding that a development-specific condensin I contributes to PDE in *Paramecium* paves the way to future investigations on non-canonical roles of SMC complexes that will further extend the repertoire of their multifaceted roles in genome biology.

## MATERIALS AND METHODS

### *Paramecium* strains and culture conditions

Unless otherwise indicated, *Paramecium tetraurelia* strain 51^86^ or its mutant variant 51 *nd7-1*^31,87^ were grown at 27°C in a wheat grass powder (WGP, Pines International) infusion medium inoculated with *Klebsiella pneumoniae* (Kp medium) or *Escherichia coli* HT115 bacteria^88^ the day before use, and supplemented with 0.8 μg/ml β-sitosterol and, when appropriate, 100 μg/mL ampicillin. Cultivation and autogamy were carried out as described^89,90^. Following starvation, the progression of autogamy was monitored cytologically by DAPI or Hoechst-staining of fixed cells. All time-points refer to hours after the T0 time-point, defined as the time when 50% of cells in the population exhibit a fragmented maternal MAC.

### RNAi-mediated gene knockdown (KD) during autogamy

Double-stranded RNA (dsRNA)-induced RNAi was carried out using the “feeding” procedure^91,92^. Two related protocols were used indifferently. In the standard protocol, *Paramecium* cells grown in Kp medium for at least 20 divisions were washed in plasmid-containing HT115 induced for dsRNA production and grown for 6 to 8 additional vegetative divisions before the onset of autogamy. In a slightly modified protocol^30^, *Paramecium* cells were grown for 10 to 15 vegetative divisions in the presence of plasmid-free HT115, then transferred to non-induced plasmid-containing HT115 medium for ∼4 divisions. They were finally fed with induced plasmid-containing HT115 for ∼8 additional divisions until the onset of autogamy. In all experiments, a survival assay was performed on 30 individual autogamous cells two or three days after T0 to test the ability of their respective progeny to form a functional new MAC (results of all survival assays can be found in Supplementary File S1).

*PGM*^29^ and *PGML*^30^ KDs were performed using published RNAi-inducing plasmids. New constructs were made to knock down *SMC4.1* (p0424), *SMC4.2* (p0421) and *CAPD2.3* (p0428, p0429; p0428 was used for all KDs shown except for two replicates used for the analysis of post-autogamous progeny survival). For the simultaneous KD of *CAPH.3a* and *CAPH.3b*, cultures of HT115 carrying either pMG16 or pMG17 (localization of CapH.3-FLAG and immunostaining of histone marks), or p0456 or p0457 (other experiments) were mixed in equivalent amounts before feeding *Paramecium* cells. Double KDs of duplicated genes originating from the last whole genome duplication^64^ are referred to collectively (e.g. *CAPH.3* KD for *CAPH.3a+b* KD). Empty vector L4440^93^ or plasmids targeting the non-essential *ND7*^87,94^ or *ICL7a*^95,96^ genes were used as controls. All plasmid maps generated in this study are available in Zenodo.

### Transgene microinjection

C-terminal fusions to the TurboID (TbID) hyperactive protein biotin ligase^55^ were expressed from plasmids p0380 or p0381 (Pgm-TbID), p0398 (PgmL4a-FLAG-TbID) or p0400 (PgmL4a-TbID). Plasmids p0460, p0475, p0476, p0464 and pTB23 were used to express CapD2.3-HA, CapD2.3-FLAG-HA, Smc4.1-FLAG, Smc4.2-FLAG and CapH.3a-FLAG, respectively. All transgenes, engineered to resist RNAi against their corresponding endogenous genes, were expressed under the control of their respective endogenous regulatory sequences.

Microinjection of linearized plasmid DNA was performed as described^97^, following restriction by BsaAI, AlwNI or XmnI (New England Biolabs). For TbID proximity labeling and the localization of CapD2.3-HA, Smc4.1-Flag and Smc4.2-Flag fusions, linearized transgene-carrying plasmids were co-injected into the MAC of vegetative *P. tetraurelia* 51 *nd7-1* with a 1:6 (TbID experiments) or 1:10 mass ratio of a linearized *ND7*-complementing plasmid, as described^31^. Transformants were selected on their trichocyst discharge ability, and transgene injection levels were estimated by qPCR^59^, using either the *KU80c* (PTET.51.1.G1140146)^35^ or the *51A* surface antigen gene (PTET.51.1.G1060180)^86^ as genomic references (see Zenodo). All qPCR primer pairs are listed in Table S3. For CapD2.3-FLAG-HA immunoprecipitation experiments and the localization of CapH.3-FLAG, linearized plasmids were solo-injected into the MAC of vegetative *P. tetraurelia* 51. Transformants were selected using transgene-specific PCR primers.

### Protein proximity labeling by *in vivo* TbID-mediated biotinylation

#### Proximity labeling during autogamy

Individual transformants harboring RNAi-resistant plasmids p0381 (*PGM*-TbID*) or p0398 (*PGML4a*- FLAG-TbID*) and non-injected control clones were grown in BHB wheatgrass infusion^89^, or in WGP when indicated, and subjected to control *ND7* KD in 5 to 10-mL cultures, and to *PGM* or *PGML4* KD, respectively, in 2-L cultures. At T5 during autogamy, 1-mL aliquots of each culture were kept at 27°C to proceed with survival assays. For *PGM* and *PGML4* KDs, biotin powder (Sigma Aldrich) was directly added to the rest of the cultures (500 µM final concentration). After a 1-hour incubation with biotin at 27°C, 1-mL aliquots of the cultures were stored at 27°C for three more days for survival assays. The remaining cells were collected by centrifugation and washed three times in 100 mL Dryl buffer (2 mM sodium citrate, 1 mM NaH_2_PO_4_, 1 mM Na_2_HPO_4_, 1 mM CaCl_2_). Cell pellets were snap-frozen in liquid N_2_ and stored at -80°C.

To verify the functionality of the Pgm-TbID and PgmL4-FLAG-TbID fusion proteins, survival assays were performed at T∼72 during autogamy (day 4, see Figure S1) on the progeny of the transformed and non-injected clones stored at 27°C before or after biotin addition (Supplementary File S1).

#### Recovery of biotinylated proteins

Frozen cell pellets were lysed by direct addition of 1.6-volume of boiling lysis buffer A (4% SDS, 4X cOmplete ULTRA Roche protease inhibitor, 1 mM AEBSF) and incubating the lysate at 95°C for 10 minutes (min) with frequent vortexing. Following centrifugation (21,500g for 10 min at room temperature), the lysate was diluted by adding 2 volumes of binding buffer (50 mM Tris pH7.4, 150 mM NaCl, 5 mM EDTA, 2% Triton) completed with protease inhibitors (final concentrations: 4X cOmplete ULTRA Roche protease inhibitor, 1 mM 4-(2-Aminoethyl) benzenesulfonyl fluoride (AEBSF, Sigma Aldrich)) and dialyzed overnight in Spectra/Por 12-14,000 MW membrane tubes at 4°C in 1 L of dialysis buffer (50 mM Tris pH 7.4, 150 mM NaCl, 5 mM EDTA, 0,4% Triton, 0,1% SDS) to eliminate residual free biotin. The dialyzed lysate was added to 200 µL of Pierce^TM^ Streptavidin agarose beads (Thermo Scientific) pre-equilibrated with binding buffer, and incubated overnight at 4°C on a rotating wheel. The streptavidin-bound fraction was collected by centrifugation (∼2400g for 5 min at 4°C) and washed 3 times in 1 mL of wash buffer (50 mM Tris pH 7.4, 500 mM NaCl, 5 mM EDTA, 0.2% SDS, 2% Triton, 4X cOmplete ULTRA Roche protease inhibitor, 1 mM AEBSF), followed by 3 washes in 1 mL of TAP buffer (10% glycerol, 0.1% NP40, 2 mM EDTA pH8, 50 mM HEPES pH 7.9, 100 mM KCl). Streptavidin agarose beads were resuspended in 50 µL H_2_O + 4X cOmplete ULTRA Roche protease inhibitor + 1 mM AEBSF + 50 µL Laemmli buffer 4X + 200 mM DTT and incubated at 98°C for 10 min. For silver staining (Silver Stain Plus kit Bio-Rad) and western blot analysis, 7.5 µL of beads were loaded on 4-15% Mini-PROTEAN® TGX™ precast polyacrylamide gels (Bio Rad) and run in Tris-glycine buffer as described^31^. For liquid chromatography coupled to tandem mass spectrometry (LC-MS/MS), 45 µL of each streptavidin-bound fraction were loaded onto a 10-well 4-15% Mini-PROTEAN TGX stain-free protein gel (Bio Rad) and run for 15 min at 120 V in Tris-glycine buffer. The gel was stored at 4°C in ultrapure water until use.

### Serial nuclear extraction

Nuclear extracts prepared at increasing salt concentrations (15 mM; 150 mM; 300 mM NaCl) from cells expressing the *CAPD2.3-FLAG-HA* transgene were performed as previously described^46^. 10^6^ autogamous cells (T=∼10 hours) were lysed with a Potter-Elvehjem homogenizer in 3 volumes of lysis buffer B (10 mM Tris pH 6.8, 10 mM MgCl_2_, 0.2% Nonidet P-40, 1 mM PMSF, 4 mM benzamidine, 1x Complete EDTA-free Protease Inhibitor Cocktail tablets (Roche)). The nuclei-containing pellet was collected by centrifugation and washed with the addition of 2.5 volumes of washing solution A (0.25 M sucrose, 10 mM MgCl_2_, 10 mM Tris pH 7.4, 1 mM PMSF, 4 mM benzamidine, 1x Complete EDTA-free Protease Inhibitor Cocktail tablets (Roche)). The pellet was incubated in 1 volume of 2X nuclear extraction buffer A (100 mM Hepes pH 7.8, 100 mM KCl, 30 mM NaCl, 0.2 mM EDTA, 20% Glycerol, 2 mM DTT, 0.02% Nonidet P-40, 2 mM PMSF, 2x Complete EDTA-free Protease Inhibitor Cocktail tablets (Roche)) for 1 hour at 4°C. The salt-extractable fraction at 15 mM NaCl was recovered following centrifugation for 3 min at 10,000 g at 4°C. The insoluble fraction was resuspended in 2X nuclear extraction buffer A (same as above except for 300 mM NaCl) and incubated for 1h at 4°C. The salt-extractable fraction at 150 mM NaCl was recovered following centrifugation for 3 min at 10,000 g at 4°C. The insoluble fraction was resuspended in 2X nuclear extraction buffer A (same as above except for 600 mM NaCl) and incubated for 1h at 4°C. The salt-extractable fraction at 300 mM NaCl was recovered following centrifugation for 3 min at 10,000 g at 4°C.

### Immunoprecipitation

#### Immunoprecipitation of CapD2.3-FLAG-HA

*Paramecium* nuclear protein extracts were prepared as previously described^46^. 3x10^6^ autogamous cells (T=∼10 hours) were lysed with a Potter-Elvehjem homogenizer in 3 volumes of lysis buffer B. The nuclei-containing pellet was collected by centrifugation and washed with the addition of 2.5 volumes of washing solution. The pellet was incubated in 1 volume of 2X nuclear extraction buffer containing 300 mM NaCl for 1 hour at 4°C. The salt-extractable fraction at 150 mM NaCl was recovered following centrifugation for 3 minutes at 10,000 g at 4°C. Nuclear extracts were incubated overnight at 4°C with 150 μl anti-FLAG M2 magnetic beads (M8823, Sigma) pre-washed with 1 mL TEGN buffer (20 mM Tris pH 8, 0.1 mM EDTA, 10% Glycerol, 150 mM NaCl, 0.01% Nonidet P-40). Beads were washed five times with TEGN buffer.

#### Tandem affinity immunoprecipitation of CapD2.3-FLAG-HA

FLAG IP was performed as described above. The beads were eluted with 3xFLAG peptide (F4799, Sigma-Aldrich) and the same volume of TEGN buffer at 4°C for 5 hours. The FLAG eluate was further affinity-purified on anti-HA antibody-conjugated agarose (A2095, Merck) overnight at 4°C and eluted with the HA peptide (I21149, Merck) overnight at 4°C.

### Mass spectrometry

#### LC MS/MS of proteins bound to streptavidin agarose beads

##### Sample preparation

Gel plugs were destained in a solution of ACN/NH_4_HCO_3_ 50mM (50/50) for 15 minutes with agitation. Plugs were reduced in 10 mM DTT for 45 min at 56°C, then alkylated in 55 mM IAA for 45 min at room temperature. After a wash and dehydration step, proteins in the plugs were trypsinized overnight at 37°C in a 25 mM NH_4_HCO_3_ buffer (0.2 µg high sequencing grade trypsin from Promega in 20 µL buffer). The resulting peptides were desalted using C18 ZipTip (Pierce Biotechnology) prior to injection.

##### LC-MS/MS

Peptides were analyzed using a Q-Exactive Plus coupled to a Nano-LC Proxeon 1000, both from ThermoFisher Scientific (Waltham, MA, USA). Peptides were separated by chromatography with the following settings: Acclaim PepMap100 C18 pre-column (0.075 x 20 mm, 3 μm, 100 Å), Pepmap-RSLC Proxeon C18 column (0.075 x 500 mm, 2 μm, 100 Å), 300 nl/min flow rate, a 98-min gradient from 95% solvent A (H_2_O/0.1% FA) to 35 % solvent B (100 % ACN/0.1% FA) followed by column regeneration, giving a total time of 120 minutes. Peptides were analyzed in the Orbitrap cell in positive mode, at a resolution of 70,000, with a mass range of m/z 375-1500 and an AGC target of 3.10^6^. MS/MS data were acquired in the Orbitrap cell in a Top20 mode. Peptides were selected for fragmentation by Higher-energy C-trap Dissociation (HCD) with a Normalized Collisional Energy of 27%, a dynamic exclusion of 60 seconds, and a quadrupole isolation window of 1.4 Da and an AGC target of 2.10^5^. Peptides with unassigned charge states or monocharged peptides were excluded from the MS/MS acquisition. The maximum ion accumulation times were set to 50 ms for MS acquisition and 45 ms for MS/MS acquisition

##### Data analysis

MS raw files were processed using PEAKS Online X (build 1.6, Bioinformatics Solutions Inc.). The data was searched against the *Paramecium tetraurelia* protein database (last modified in 2017) available from ParameciumDB (https://paramecium.i2bc.paris-saclay.fr/download/Paramecium/tetraurelia/51), consisting of 40,460 entries and further modified with the addition of 6 sequences corresponding to the two bait constructs, the birA and TurboID biotin ligases^55,58^, streptavidin and the FLAG peptide. Parent mass tolerance was set to 5 ppm and fragment mass tolerance to 0.05 Da. Specific tryptic cleavage was selected and a maximum of 2 missed cleavages was authorized. For identification, the following post-translational modifications were included: oxidation (M), deamidation (NQ) as variables and carbamidomethylation (C) as fixed. The maximum number of variable modifications per peptide was limited to 2. Peptide identifications were filtered based on a 1% FDR (False Discovery Rate) threshold at PSM level. Match between runs was performed with a mass error tolerance of 5 ppm, a retention time shift tolerance of 1 min and a feature intensity threshold of 100. A normalization between runs was applied based on the Total Ion Current (TIC) and protein abundance was finally inferred using a Top N label free quantification approach. Multivariate statistics on protein measurements and graphical results were generated using Qlucore Omics Explorer 3.8.9 (Qlucore AB, Lund, SWEDEN). A positive threshold value of 1 was specified to enable a log2 transformation of abundance data for normalization *i.e.* all abundance data values below the threshold were replaced by 1 before transformation. The transformed data were finally used for statistical comparison of groups using a Student’s bilateral t-test.

Protein identification was performed for all samples together and led to the identification of 1471 protein groups (see Zenodo). Two samples (Pgm-1 and CTL-8) were considered as outliers because of their very low material abundance and final protein content identification, and were finally excluded from the statistical analysis.

#### LC MS/MS of protein immunoprecipitates

##### Sample preparation

Beads from pulldown experiments were incubated overnight at 37°C with 40 μL of 50 mM NH_4_HCO_3_ (Sigma-Aldrich) buffer containing 1 µg of sequencing-grade trypsin/Lys C mix (Promega). The digested peptides were loaded and desalted on evotips (Evosep, Odense, Denmark) according to the manufacturer’s procedure before LC-MS/MS analysis.

##### LC-MS/MS acquisition

Samples were analyzed on a timsTOF Pro 2 mass spectrometer (Bruker Daltonics, Bremen, Germany) coupled to an Evosep one system (Evosep, Odense, Denmark) operating with the 30SPD method developed by the manufacturer. Briefly, the method is based on a 44-min gradient and a total cycle time of 48 min with a C18 analytical column (0.15 x 150 mm, 1.9 µm beads, ref EV-1106) equilibrated at 40°C and operated at a flow rate of 500 nL/min. H_2_O/0.1 % FA was used as solvent A and ACN/ 0.1 % FA as solvent B. For the tandem affinity purification of CapD2.3, the timsTOF Pro 2 was operated with a DIA-PASEF method comprising 12 pydiAID frames with 3 mass windows per frame resulting in a cycle time of 0.975 seconds, as described in Bruker application note LCMS 218. For the FLAG affinity purification, the timsTOF Pro 2 was operated in DDA-PASEF mode over a 1.3 sec cycle time at 60 Hz MS/MS scanning rate. Mass spectra for MS and MS/MS scans were recorded between 100 and 1700 m/z.

##### Data analysis

For the tandem affinity purification of CapD2.3-FLAG-HA, MS raw files were processed using Spectronaut 18 (Biognosys, Switzerland). Data were searched against the same *Paramecium* protein database as above, combined with the SwissProt *Mus musculus* database (12-2023, 17799 entries). Contaminating proteins originating from the anti-FLAG antibody-coupled beads used for IP and identified only in the *Mus musculus* taxonomy were excluded from the final protein list. Specific tryptic cleavages were selected and a maximum of 2 missed cleavages were allowed. The following post-translational modifications were considered for identification: Acetyl (Protein N-term) and Oxidation (M) as variable and Cys-Cys (C) as fixed. The maximum number of variable modifications was set to 5. Identifications were filtered based on a 1% precursor and protein Qvalue cutoff threshold. The protein LFQ method was set to automatic and the quantity was set at the MS2 level with a cross-run normalization applied. Multivariate statistics on protein measurements were performed as described above, using Qlucore Omics Explorer 3.9 (Qlucore AB, Lund, SWEDEN).

For the affinity purification of CapD2.3, MS raw files were processed using PEAKS Online 11 (build 1.9, Bioinformatics Solutions Inc.). Data were searched against the same *Paramecium* protein database as above, with PTET.51.1.P0430205 sequence modified to include the HA tag of the bait. Parent mass tolerance was set to 20 ppm, with fragment mass tolerance of 0.05 Da. Semi-specific tryptic cleavages was selected and a maximum of 1 missed cleavage was allowed. The following post-translational modifications were considered for identification: Oxidation (M), Deamidation (NQ) and Acetylation (Protein N-term) as variable and Half of a disulfide bridge (C) as fixed. Identifications were filtered to a <1% FDR (False Discovery Rate) threshold at both peptide and protein group levels. Label free quantification was performed using the PEAKS Online 11 quantification module, allowing a mass tolerance of 5 ppm, a CCS error tolerance of 0.01 and a 0.5-min retention time shift tolerance for match between runs. Protein abundance was inferred using the top N peptide method and TIC was used for normalization. Multivariate statistics, data normalization and differential statistical analysis were performed as described for the tandem affinity purification experiment.

### Immunofluorescence and quantification

#### Detection of biotinylated proteins, Smc4.1-FLAG, Smc4.2-FLAG, CapD2.3-HA, Pgm and PgmL proteins

Cells were first permeabilized in 1% Triton X-100, then fixed in 2% formaldehyde and processed for immunofluorescence labeling and epifluorescence microscopy as described^30^. Biotinylated proteins were detected using Alexafluor (AF) 488-conjugated Streptavidin (S11223, Thermo Fisher Scientific). FLAG- and HA-tagged proteins were detected using anti-FLAG M2 (F1804, Sigma Aldrich) or anti-HA (H9658, Sigma Aldrich) mouse monoclonal antibodies. Pgm, PgmL1 and PgmL5 were detected using guinea pig anti-Pgm 2659-GP2^31^ & GP3 (obtained as described for GP2 from a different animal, see Figure S8), guinea pig anti-PgmL1^51^ and rabbit anti-PgmL5^30^ polyclonal antibodies, respectively. Anti-PgmL2, anti-PgmL3 and anti-PgmL4 antibodies were obtained following guinea pig immunization with C-terminally His-tagged peptides PgmL2[aa 1-285], PgmL3a[aa 438-735] and PgmL4a[aa 2-177], respectively, followed by antigen affinity purification (Proteogenix). Their specificity was confirmed by whole-cell immunofluorescence labeling upon the KD of their respective endogenous target PgmL (Figure S8). Primary antibodies were revealed with AF 488 or 568-conjugated anti-rabbit, anti-mouse or anti-guinea pig secondary antibodies (Thermo Fisher Scientific) as described^30^. All antibodies and their working dilutions are listed in Table S4. The intensity of the immunofluorescence signal in the developing new MACs was quantified using ImageJ as described^30^. Nuclei were selected on their size (in µm^2^ at the maximal area section), in order to compare populations within the size range corresponding to the peak of signal intensities in the control. Fluorescence intensities were normalized against the median intensity of each respective control (see Zenodo).

#### Detection of CapH.3-FLAG, CapD2.3-FLAG-HA, H3K9me3 and H3K27me3

Cells were fixed for 30 min in solution I (10 mM EGTA, 25 mM HEPES, 2 mM MgCl_2_, 60 mM PIPES pH 6.9, 1X PHEM; 1% formaldehyde, 2.5% Triton X-100, 4% sucrose), and for 10 min in solution II (1X PHEM, 4% formaldehyde, 1.2% Triton X-100, 4% sucrose). Following blocking in 3% bovine serum albumin-supplemented Tris buffered saline-Tween 20 0.1% (TBST) for 10 min, fixed cells were incubated overnight at room temperature under agitation with the following primary antibodies: rabbit anti-H3K9me3 and anti-H3K27me3^46^, guinea pig anti-Pgm 2659-GP2 and mouse anti-FLAG (MAI-91878, Thermo Fisher Scientific). Cells were labelled with AF 568-conjugated goat anti-rabbit IgG, AF 488-conjugated goat anti-guinea pig IgG or AF 568-conjugated goat anti-mouse IgG at 1:500 for 1 hour, stained with 1 μg/mL Hoechst for 5–10 min and finally mounted in Citifluor AF2 glycerol solution (antibodies and working dilutions in Table S4).

Images were acquired using a Zeiss LSM 980 laser-scanning confocal microscope with an Airyscan 2 post--processing treatment and a Plan-Apochromat 63 × /1.40 oil DIC M27 or a Plan-Apochromat 40 × /1.3 oil DIC objective. Z-series were performed with Z-steps of 0.25 μm. Images of CapH.3-FLAG localization were acquired on a Zeiss Axio Observer with a 40 X Plan Apo ON 1.3 Oil DIC (40-1). Quantification was performed as previously described^98^ using ImageJ. The volume of the nucleus (in voxels) was estimated as follows: using the Hoechst channel, the top and bottom Z stacks of the developing MAC were defined to estimate nucleus height in pixels. The equatorial Z stack of the developing MAC was defined, and the corresponding developing MAC surface was measured in pixels. The estimated volume of the developing MAC was then calculated as the product of the obtained nucleus height by the median surface. For each Z stack of the developing MAC, the H3K9me3, H3K27me3 or FLAG fluorescence intensity was measured and corrected using the ImageJ “subtract background” tool. The sum of the corrected H3K9me3, H3K27me3 or FLAG fluorescence intensities for all the Z stacks, which corresponds to the total H3K9me3, H3K27me3 or FLAG fluorescence intensity, was divided by the estimated volume to obtain the H3K9me3, H3K27me3 or FLAG fluorescence intensity per voxel in each nucleus. For foci quantification of H3K9me3 and H3K27me3 signals in the developing MACs, the standard deviation of the fluorescence of H3K9me3, H3K27me3 signals was measured with ImageJ for each Z stack and a mean standard deviation was calculated. For each condition, at least 25 nuclei were quantified. Mann-Whitney statistical tests were performed with GraphPad Prism.

### Western blot

For western blot, sample preparation and gel electrophoresis were carried out according to standard procedures. For the preparation of whole-cell extracts, around 50 mL of cells were collected and lyzed in boiling 10% SDS as described^31,98^. Samples were loaded onto 10% or 4-15% polyacrylamide Mini-PROTEAN® TGX™ Precast Protein Gels (BIO-RAD) and run in Tris-glycine migration buffer (25 mM Tris-base, 200 mM glycine, 0.1% SDS; specific information in Zenodo for each gel). Unless otherwise indicated, proteins were electro-transferred to 0.45 µm NC Protran nicrocellulose blotting membranes (Amersham) in 20 mM NaH_2_PO_4_, 20 mM Na_2_HPO_4_, pH 6.7. Biotinylated proteins were detected using horseradish peroxidase (HRP)-conjugated Streptavidin (Sigma) and enhanced chemiluminescence (ECL).

For the detection of CapD2.3-HA, Smc4.2-FLAG, Pgm and PgmL1 to PgmL4, the same antibodies were used as for immunofluorescence labeling, with the exception of anti-PgmL1, anti-PgmL2 and anti-PgmL3 antibodies, which were obtained from different animals (Proteogenix, see Table S4). The anti α-tubulin mouse monoclonal antibody TEU435^99^ was used for loading controls. Primary antibodies were revealed by ECL using species-appropriate HRP-conjugated secondary antibodies: anti-mouse IgG (Promega) or anti-guinea pig IgG (Thermo Scientific).

For the detection of CapH.3-FLAG and CapD2.3-FLAG-HA, blotting was performed overnight using a nitrocellulose membrane (GE10600002, Merck). Anti-FLAG (MAI-91878, Thermo Fisher Scientific), anti-*Paramecium* H3^46^, anti α-tubulin (05-829, Merck) or anti-HA (H6908, Merck) were used as primary antibodies. Secondary HRP-conjugated anti-mouse or anti-rabbit IgG antibodies (Promega) were used at 1:2,500 or 1: 8,000 dilution followed by ECL detection, or fluorescent secondary anti-rabbit antibodies (Biorad) (antibodies and working dilutions in Table S4).

### Nuclei purification, whole-genome sequencing and data analysis

Developing new MACs upon *PGM*, *SMC4.2*, *CAPH.3*, *CAPD2.3* or control KD were obtained from 800 mL of autogamous cells harvested at T30, immunostained using anti-PgmL1 antibodies (Table S4) and purified by fluorescence-activated nuclear sorting as described^51^. New MAC DNA was extracted using the QIAamp DNA micro kit (Qiagen) and 200 ng were fragmented with a S220 focused-ultrasonicator (Covaris) as described^51^. Sequencing libraries were prepared using the NEBNext Ultra II DNA library Prep Kit (New England Biolabs), purified on AMPure XP beads (Beckman Coulter). Following quality control (TapeStation or Bioanalyzer, Agilent) libraries were processed for paired-end sequencing using Illumina NextSeq sequencers (58 to 106M reads) at the I2BC next-generation sequencing facility or at Novogene (paired-end read length 2 x ∼75-nt or 2 x ∼150-nt, respectively). Sequencing data were filtered on expected contaminants (ribosomal DNA, mitochondrial and bacterial genomes), before mapping on the *P. tetraurelia* MAC (ptetraurelia_mac_51.fa), MAC+IES (ptetraurelia_mac_51_with_ies.fa) and MIC (ptetraurelia_mic2.fa) reference genomes using Bowtie2 (v2.2.9 --local -X500)^20,22^. Sequencing metrics are provided in Table S2. For IES analysis, the v1 annotation (internal_eliminated_sequence_PGM_ParTIES.pt_51.gff3) was used and IES retention scores were calculated with the MIRET module of the ParTIES suite (v1.06, default parameters; https://github.com/oarnaiz/ParTIES)^100^. For TE coverage analysis, only transposable elements (TEs) from annotation v1.0 (ptetraurelia_mic2_TE_annotation_v1.0.gff3), with lengths exceeding 500 nt on contigs longer than 2 kb, were considered. Normalized TE coverage, expressed as reads per kilobase per million mapped reads (RPKM), was determined from read counts using htseq-count (v0.11.2, -- mode = intersection-nonempty; for RNA-seq, --stranded = yes)^101^ on filtered BAM files (SAMtools v1.3.1 -q 10), then normalized by the total number of reads mapped to the MIC genome. Graphics were generated with R (v4.0.4) using the following packages: *ComplexHeatmap* (v2.6.2)^102^, *rtracklayer*(v1.50) ^103^ and *circlize* (v0.4.13)^104^.

## DATA AVAILABILITY

The accession numbers of all condensin subunit genes are listed in Supplementary Table S1 and refer to the ParameciumDB database (https://paramecium.i2bc.paris-saclay.fr/)^66^. Plasmid maps and nucleotide sequences, raw images (microscopy, western blots), statistical data, scripts and numerical data underlying this article can be found in the Zenodo open-access research data repository (https://zenodo.org/records/17255484). The sequencing data underlying this article are available in the European Nucleotide Archive (ENA, https://www.ebi.ac.uk/ena/browser/home)^105^ under accession number PRJEB96275. The mass spectrometry proteomics data have been deposited to the ProteomeXchange Consortium via the PRIDE partner repository (https://www.ebi.ac.uk/pride/)^106^ with the dataset identifier PXD068119.

## ACKNOWLEDGEMENTS

We acknowledge the ImagoSeine facility for confocal miscroscopy, the ProteoSeine facility of Institut Jacques Monod for mass spectromery, the sequencing and bioinformatics expertise of the I2BC High-throughput sequencing facility, supported by *France Génomique* (funded by the French National Program *Investissement d’Avenir* ANR-10-INBS-09). The present work has benefited from the Imagerie-Gif core facility supported by the *Agence Nationale de la Recherche* (ANR-10-INBS-04 / FranceBioImaging; ANR-11-IDEX-0003-02 / Saclay Plant Sciences). We are grateful to Linda Sperling for critical reading of the manuscript and to Estienne Swart and Aditi Singh for sharing their data before publication. We thank Pascaline Tirand and Fanny Culot for their assistance in maintaining the I2BC *Paramecium* stock collection, and Vincent Maupu-Massamba, Nelly Sainsard and Cristina Delawarde for medium preparation.

This work was funded by the *Centre National de la Recherche Scientifique* (CNRS); the *Agence Nationale pour la Recherche* (ANR) [project “POLYCHROME” ANR-19-CE12-0015 to SD and OA], [project “CURE” ANR-21-CE12-0019 to MB], [project “SELECTION” ANR-23-CE12-0027 to SD and GC], [project “MODIFICATION” ANR-25-CE12-7757 to SD]; the *Fondation pour la Recherche Médicale* [“Equipe FRM EQU202103012766” to MB and “Equipe FRM EQU202203014643” to SD]; the French Infrastructure for Integrated Structural Biology (FRISBI) (ANR-10-INBS-0005). TB was recipient of doctoral fellowships from BioSPC Doctoral School from Université Paris Cité and *Fondation pour la Recherche Médicale* (FDT202404018139) and a post-doc transition fellowship from EUR GENE [ANR-17-EURE-0013].

## REFERENCES

1. Cosby, R. L., Chang, N. C. & Feschotte, C. Host-transposon interactions: conflict, cooperation, and cooption. Genes Dev 33, 1098–1116 (2019).

2. Darmon, E. & Leach, D. R. F. Bacterial Genome Instability. Microbiol. Mol. Biol. Rev. 78, 1–39 (2014).

3. Koonin, E. V., Makarova, K. S., Wolf, Y. I. & Krupovic, M. Evolutionary entanglement of mobile genetic elements and host defence systems: guns for hire. Nat. Rev. Genet. 21, 119–131 (2020).

4. Vasu, K. & Nagaraja, V. Diverse Functions of Restriction-Modification Systems in Addition to Cellular Defense. Microbiol. Mol. Biol. Rev. 77, 53–72 (2013).

5. Hille, F. et al. The Biology of CRISPR-Cas: Backward and Forward. Cell 172, 1239–1259 (2018).

6. Ishino, Y., Krupovic, M. & Forterre, P. History of CRISPR-Cas from Encounter with a Mysterious Repeated Sequence to Genome Editing Technology. J. Bacteriol. 200, 10.1128/jb.00580-17 (2018).

7. Deep, A. et al. The SMC-family Wadjet complex protects bacteria from plasmid transformation by recognition and cleavage of closed-circular DNA. Mol. Cell 82, 4145–4159.e7 (2022).

8. Liu, H. W. et al. DNA-measuring Wadjet SMC ATPases restrict smaller circular plasmids by DNA cleavage. Mol. Cell 82, 4727–4740.e6 (2022).

9. Errington, J. Regulation of endospore formation in Bacillus subtilis. Nat. Rev. Microbiol. 1, 117–126 (2003).

10. Kunkel, B., Losick, R. & Stragier, P. The Bacillus subtilis gene for the development transcription factor sigma K is generated by excision of a dispensable DNA element containing a sporulation recombinase gene. Genes Dev 4, 525–535. (1990).

11. Kapitonov, V. V. & Jurka, J. RAG1 core and V(D)J recombination signal sequences were derived from Transib transposons. PLoS Biol 3, e181. Epub 2005 May 24. (2005).

12. Liu, C., Zhang, Y., Liu, C. C. & Schatz, D. G. Structural insights into the evolution of the RAG recombinase. Nat. Rev. Immunol. 22, 353–370 (2022).

13. Dedukh, D. & Krasikova, A. Delete and survive: strategies of programmed genetic material elimination in eukaryotes. Biol. Rev. 97, 195–216 (2022).

14. Drotos, K. H. I., Zagoskin, M. V., Kess, T., Gregory, T. R. & Wyngaard, G. A. Throwing away DNA: programmed downsizing in somatic nuclei. Trends Genet. TIG 38, 483–500 (2022).

15. Mochizuki, K. Programmed DNA elimination. Curr. Biol. 34, R843–R847 (2024).

16. Wang, J. & Davis, R. E. Programmed DNA elimination in multicellular organisms. Curr Opin Genet Dev 27, 26–34 (2014).

17. Estrem, B., Davis, R. E. & Wang, J. End resection and telomere healing of DNA double-strand breaks during nematode programmed DNA elimination. Nucleic Acids Res. 52, 8913–8929 (2024).

18. Rey, C., Launay, C., Wenger, E. & Delattre, M. Programmed DNA elimination in Mesorhabditis nematodes. Curr. Biol. 33, 3711–3721.e5 (2023).

19. Prescott, D. M. The DNA of ciliated protozoa. Microbiol Rev 58, 233–267 (1994).

20. Arnaiz, O. et al. The Paramecium germline genome provides a niche for intragenic parasitic DNA: evolutionary dynamics of internal eliminated sequences. PLoS Genet. 8, e1002984 (2012).

21. Chen, X. et al. The architecture of a scrambled genome reveals massive levels of genomic rearrangement during development. Cell 158, 1187–1198 (2014).

22. Guérin, F. et al. Flow cytometry sorting of nuclei enables the first global characterization of Paramecium germline DNA and transposable elements. BMC Genomics 18, 327 (2017).

23. Hamilton, E. P. et al. Structure of the germline genome of Tetrahymena thermophila and relationship to the massively rearranged somatic genome. eLife 5, e19090 (2016).

24. Sellis, D. et al. Massive colonization of protein-coding exons by selfish genetic elements in Paramecium germline genomes. PLoS Biol. 19, e3001309 (2021).

25. Bétermier, M., Klobutcher, L. A. & Orias, E. Programmed chromosome fragmentation in ciliated protozoa: multiple means to chromosome ends. Microbiol. Mol. Biol. Rev. MMBR 87, e0018422 (2023).

26. Bétermier, M. & Duharcourt, S. Programmed Rearrangement in Ciliates: Paramecium. Microbiol. Spectr. 2, (2014).

27. Gratias, A. & Bétermier, M. Processing of double-strand breaks is involved in the precise excision of paramecium internal eliminated sequences. Mol. Cell. Biol. 23, 7152–7162 (2003).

28. Balan, T., Lerner, L. K., Holoch, D. & Duharcourt, S. Small-RNA-guided histone modifications and somatic genome elimination in ciliates. WIREs RNA 15, e1848 (2024).

29. Baudry, C. et al. PiggyMac, a domesticated piggyBac transposase involved in programmed genome rearrangements in the ciliate Paramecium tetraurelia. Genes Dev. 23, 2478–2483 (2009).

30. Bischerour, J. et al. Six domesticated PiggyBac transposases together carry out programmed DNA elimination in Paramecium. eLife 7, e37927 (2018).

31. Dubois, E. et al. Multimerization properties of PiggyMac, a domesticated piggyBac transposase involved in programmed genome rearrangements. Nucleic Acids Res 45, 3204–3216 (2017).

32. Singh, M. et al. Origins of genome-editing excisases as illuminated by the somatic genome of the ciliate Blepharisma. Proc. Natl. Acad. Sci. U. S. A. 120, e2213887120 (2023).

33. Abello, A. et al. Functional diversification of Paramecium Ku80 paralogs safeguards genome integrity during precise programmed DNA elimination. PLoS Genet. 16, e1008723 (2020).

34. Kapusta, A. et al. Highly precise and developmentally programmed genome assembly in Paramecium requires ligase IV-dependent end joining. PLoS Genet. 7, e1002049 (2011).

35. Marmignon, A. et al. Ku-mediated coupling of DNA cleavage and repair during programmed genome rearrangements in the ciliate Paramecium tetraurelia. PLoS Genet. 10, e1004552 (2014).

36. Verron, B. et al. The linker region of a development-specific DNA polymerase X ensures efficient repair of programmed DNA double-strand breaks in Paramecium tetraurelia. Nucleic Acids Res. 53, gkaf286 (2025).

37. Bétermier, M., Borde, V. & de Villartay, J.-P. Coupling DNA Damage and Repair: an Essential Safeguard during Programmed DNA Double-Strand Breaks? Trends Cell Biol. 30, 87–96 (2020).

38. Bischerour, J. et al. Uncoupling programmed DNA cleavage and repair scrambles the Paramecium somatic genome. Cell Rep. 43, 114001 (2024).

39. Hickman, A. B. & Dyda, F. Mechanisms of DNA Transposition. Microbiol Spectr 3, MDNA3-0034–2014 (2015).

40. Morellet, N. et al. Sequence-specific DNA binding activity of the cross-brace zinc finger motif of the piggyBac transposase. Nucleic Acids Res 46, 2660–2677 (2018).

41. Klobutcher, L. A. & Herrick, G. Consensus inverted terminal repeat sequence of Paramecium IESs: resemblance to termini of Tc1-related and Euplotes Tec transposons. Nucleic Acids Res. 23, 2006–2013 (1995).

42. Charmant, O. et al. The PIWI-interacting protein Gtsf1 controls the selective degradation of small RNAs in Paramecium. Nucleic Acids Res. 53, gkae1055 (2025).

43. Lepère, G. et al. Silencing-associated and meiosis-specific small RNA pathways in Paramecium tetraurelia. Nucleic Acids Res. 37, 903–915 (2009).

44. Sandoval, P. Y., Swart, E. C., Arambasic, M. & Nowacki, M. Functional diversification of Dicer-like proteins and small RNAs required for genome sculpting. Dev Cell 28, 174–88 (2014).

45. Wang, C. et al. GTSF1 is required for transposon silencing in the unicellular eukaryote Paramecium tetraurelia. Nucleic Acids Res. 52, 13206–13223 (2024).

46. Frapporti, A. et al. The Polycomb protein Ezl1 mediates H3K9 and H3K27 methylation to repress transposable elements in Paramecium. Nat. Commun. 10, 2710 (2019).

47. Lhuillier-Akakpo, M. et al. Local effect of enhancer of zeste-like reveals cooperation of epigenetic and cis-acting determinants for zygotic genome rearrangements. PLoS Genet. 10, e1004665 (2014).

48. Miró-Pina, C. et al. Paramecium Polycomb repressive complex 2 physically interacts with the small RNA-binding PIWI protein to repress transposable elements. Dev. Cell 57, 1037–1052.e8 (2022).

49. Wang, C. et al. A small RNA-guided PRC2 complex eliminates DNA as an extreme form of transposon silencing. Cell Rep. 40, 111263 (2022).

50. Swart, E. C. et al. Identification and analysis of functional associations among natural eukaryotic genome editing components [version 1; peer review: 1 approved, 1 approved with reservations]. F1000Research 6, 1374 (2017).

51. Zangarelli, C. et al. Developmental timing of programmed DNA elimination in *Paramecium tetraurelia* recapitulates germline transposon evolutionary dynamics. Genome Res. 32, 2028–2042 (2022).

52. Bürmann, F. & Löwe, J. Structural biology of SMC complexes across the tree of life. Curr. Opin. Struct. Biol. 80, 102598 (2023).

53. Hoencamp, C. & Rowland, B. D. Genome control by SMC complexes. Nat. Rev. Mol. Cell Biol. 24, 633–650 (2023).

54. Hirano, T. Condensin-Based Chromosome Organization from Bacteria to Vertebrates. Cell 164, 847–857 (2016).

55. Branon, T. C. et al. Efficient proximity labeling in living cells and organisms with TurboID. Nat Biotechnol 36, 880–887 (2018).

56. Arnaiz, O. et al. Improved methods and resources for paramecium genomics: transcription units, gene annotation and gene expression. BMC Genomics 18, 483 (2017).

57. Kim, D. I. et al. Probing nuclear pore complex architecture with proximity-dependent biotinylation. Proc. Natl. Acad. Sci. U. S. A. 111, E2453–2461 (2014).

58. Roux, K. J., Kim, D. I., Raida, M. & Burke, B. A promiscuous biotin ligase fusion protein identifies proximal and interacting proteins in mammalian cells. J Cell Biol 196, 801–10 (2012).

59. Bazin-Gélis, M. et al. Inter-generational nuclear crosstalk links the control of gene expression to programmed genome rearrangement during the Paramecium sexual cycle. Nucleic Acids Res. 51, 12337–12351 (2023).

60. Singh, A. et al. ISWI1 complex proteins facilitate developmental genome editing in Paramecium. Genome Res. 35, 93–108 (2025).

61. Singh, A. et al. Chromatin remodeling is required for sRNA-guided DNA elimination in Paramecium. EMBO J. 41, e111839 (2022).

62. Matsuda, A. & Forney, J. D. The SUMO pathway is developmentally regulated and required for programmed DNA elimination in Paramecium tetraurelia. Eukaryot. Cell 5, 806–815 (2006).

63. Hooff, J. J. E. van, Raas, M. W. D., Tromer, E. C. & Eme, L. Repeated duplications and losses shaped SMC complex evolution from archaeal ancestors to modern eukaryotes. Cell Rep. 44, (2025).

64. Aury, J.-M. et al. Global trends of whole-genome duplications revealed by the ciliate Paramecium tetraurelia. Nature 444, 171–178 (2006).

65. Zhang, F., Bechara, S. & Nowacki, M. Structural maintenance of chromosomes (SMC) proteins are required for DNA elimination in Paramecium. Life Sci. Alliance 7, (2024).

66. Arnaiz, O., Meyer, E. & Sperling, L. ParameciumDB 2019: integrating genomic data across the genus for functional and evolutionary biology. Nucleic Acids Res. 48, D599– D605 (2020).

67. The Seventh International Meeting on Ciliate Molecular Biology Genetics Nomenclature Committee et al. Proposed Genetic Nomenclature Rules for Tetrahymena thermophila, Paramecium primaurelia and Paramecium tetraurelia. Genetics 149, 459–462 (1998).

68. Nowak, J. K. et al. Functional study of genes essential for autogamy and nuclear reorganization in Paramecium. Eukaryot Cell 10, 363–72 (2011).

69. Lees-Miller, J. P. et al. Uncovering DNA-PKcs ancient phylogeny, unique sequence motifs and insights for human disease. Prog. Biophys. Mol. Biol. 163, 87–108 (2021).

70. Ali, E. I., Loidl, J. & Howard-Till, R. A. A streamlined cohesin apparatus is sufficient for mitosis and meiosis in the protist Tetrahymena. Chromosoma 127, 421–435 (2018).

71. Howard-Till, R. & Loidl, J. Condensins promote chromosome individualization and segregation during mitosis, meiosis, and amitosis in Tetrahymena thermophila. Mol. Biol. Cell 29, 466–478 (2018).

72. Howard-Till, R., Tian, M. & Loidl, J. A specialized condensin complex participates in somatic nuclear maturation in Tetrahymena thermophila. Mol. Biol. Cell 30, 1326–1338 (2019).

73. Cervantes, M. D., Coyne, Robert S., Xi, Xiaohui & and Yao, M.-C. The Condensin Complex Is Essential for Amitotic Segregation of Bulk Chromosomes, but Not Nucleoli, in the Ciliate Tetrahymena thermophila. Mol. Cell. Biol. 26, 4690–4700 (2006).

74. Luo, Z. et al. Rearrangement of macronucleus chromosomes correspond to TAD-like structures of micronucleus chromosomes in Tetrahymena thermophila. Genome Res. 30, 406–414 (2020).

75. Villano, D. J., Prahlad, M., Singhal, A., Sanbonmatsu, K. Y. & Landweber, L. F. Widespread 3D genome reorganization precedes programmed DNA rearrangement in Oxytricha trifallax. 2024.12.31.630814 Preprint at 10.1101/2024.12.31.630814 (2025).

76. Davidson, I. F. & Peters, J.-M. Genome folding through loop extrusion by SMC complexes. Nat. Rev. Mol. Cell Biol. 22, 445–464 (2021).

77. Uhlmann, F. A unified model for cohesin function in sister chromatid cohesion and chromatin loop formation. Mol. Cell 85, 1058–1071 (2025).

78. Dai, H.-Q. et al. Loop extrusion mediates physiological Igh locus contraction for RAG scanning. Nature 590, 338–343 (2021).

79. Zhang, Y., Zhang, X., Dai, H.-Q., Hu, H. & Alt, F. W. The role of chromatin loop extrusion in antibody diversification. Nat. Rev. Immunol. 22, 550–566 (2022).

80. Lhuillier-Akakpo, M., Guérin, F., Frapporti, A. & Duharcourt, S. DNA deletion as a mechanism for developmentally programmed centromere loss. Nucleic Acids Res. 44, 1553–1565 (2016).

81. Gratias, A. et al. Developmentally programmed DNA splicing in Paramecium reveals short-distance crosstalk between DNA cleavage sites. Nucleic Acids Res 36, 3244–3251 (2008).

82. Kim, E., Gonzalez, A. M., Pradhan, B., van der Torre, J. & Dekker, C. Condensin-driven loop extrusion on supercoiled DNA. Nat. Struct. Mol. Biol. 29, 719–727 (2022).

83. Martínez-García, B. et al. Condensin pinches a short negatively supercoiled DNA loop during each round of ATP usage. EMBO J. 42, e111913 (2023).

84. Pradhan, B. et al. Loop-extrusion-mediated plasmid DNA cleavage by the bacterial SMC Wadjet complex. Mol. Cell 85, 107–116.e5 (2025).

85. Roisné-Hamelin, F., Liu, H. W., Taschner, M., Li, Y. & Gruber, S. Structural basis for plasmid restriction by SMC JET nuclease. Mol. Cell 84, 883–896.e7 (2024).

86. Preer, J. R., Preer, L. B., Rudman, B. M. & Barnett, A. J. Deviation from the universal code shown by the gene for surface protein 51A in Paramecium. Nature 314, 188–190 (1985).

87. Skouri, F. & Cohen, J. Genetic approach to regulated exocytosis using functional complementation in Paramecium: identification of the ND7 gene required for membrane fusion. Mol Biol Cell 8, 1063–71 (1997).

88. Timmons, L., Court, D. L. & Fire, A. Ingestion of bacterially expressed dsRNAs can produce specific and potent genetic interference in Caenorhabditis elegans. Gene 263, 103–112. (2001).

89. Beisson, J. et al. Mass culture of Paramecium tetraurelia. Cold Spring Harb Protoc 2010, pdb prot5362 (2010).

90. Beisson, J. et al. Maintaining clonal Paramecium tetraurelia cell lines of controlled age through daily reisolation. Cold Spring Harb Protoc 2010, pdb prot5361 (2010).

91. Beisson, J. et al. Silencing specific Paramecium tetraurelia genes by feeding double-stranded RNA. Cold Spring Harb Protoc 2010, pdb prot5363 (2010).

92. Galvani, A. & Sperling, L. RNA interference by feeding in Paramecium. Trends Genet 18, 11–12. (2002).

93. Kamath, R. S., Martinez-Campos, M., Zipperlen, P., Fraser, A. G. & Ahringer, J. Effectiveness of specific RNA-mediated interference through ingested double-stranded RNA in Caenorhabditis elegans. Genome Biol 2, RESEARCH0002. doi: 10.1186/gb-2000-2-1-research0002 (2001).

94. Garnier, O., Serrano, V., Duharcourt, S. & Meyer, E. RNA-mediated programming of developmental genome rearrangements in Paramecium tetraurelia. Mol. Cell. Biol. 24, 7370–7379 (2004).

95. Bouhouche, K., Gout, J. F., Kapusta, A., Betermier, M. & Meyer, E. Functional specialization of Piwi proteins in Paramecium tetraurelia from post-transcriptional gene silencing to genome remodelling. Nucleic Acids Res 39, 4249–64 (2011).

96. Gogendeau, D. et al. Functional diversification of centrins and cell morphological complexity. J. Cell Sci. 121, 65–74 (2008).

97. Beisson, J. et al. DNA microinjection into the macronucleus of paramecium. Cold Spring Harb Protoc 2010, pdb prot5364 (2010).

98. de Vanssay, A. et al. The Paramecium histone chaperone Spt16-1 is required for Pgm endonuclease function in programmed genome rearrangements. PLoS Genet. 16, e1008949 (2020).

99. Callen, A. M. et al. Isolation and characterization of libraries of monoclonal antibodies directed against various forms of tubulin in Paramecium. Biol Cell 81, 95–119 (1994).

100. Denby Wilkes, C., Arnaiz, O. & Sperling, L. ParTIES: a toolbox for Paramecium interspersed DNA elimination studies. Bioinformatics 32, 599–601 (2016).

101. Anders, S., Pyl, P. T. & Huber, W. HTSeq—a Python framework to work with high-throughput sequencing data. Bioinformatics 31, 166–169 (2015).

102. Gu, Z., Eils, R. & Schlesner, M. Complex heatmaps reveal patterns and correlations in multidimensional genomic data. Bioinformatics 32, 2847–2849 (2016).

103. Lawrence, M., Gentleman, R. & Carey, V. rtracklayer: an R package for interfacing with genome browsers. Bioinformatics 25, 1841–1842 (2009).

104. Gu, Z., Gu, L., Eils, R., Schlesner, M. & Brors, B. circlize implements and enhances circular visualization in R. Bioinformatics 30, 2811–2812 (2014).

105. Yuan, D. et al. The European Nucleotide Archive in 2023. Nucleic Acids Res. 52, D92–D97 (2024).

106. Perez-Riverol, Y. et al. The PRIDE database at 20 years: 2025 update. Nucleic Acids Res. 53, D543–D553 (2025).

