## Supplementary Information for "A developmental condensin I complex assists the *Paramecium* PiggyMac domesticated transposase during programmed DNA elimination"

### SUPPLEMENTARY MATERIALS AND METHODS

#### Monitoring of vegetative phenotypes

For *SMC2*, *SMC4.1* and *CAPG* KDs, the maps of all RNAi-inducing plasmids can be found in Zenodo. Two feeding inserts (Fi-1 and Fi-2) were tested independently for *CAPG* KD (Supplementary File S1).

For the survival analysis of vegetative lines, cells were grown at 27°C and counted every day to estimate the number of apparent divisions, and a single cell was picked and transferred to 200 µL of freshly induced RNAi medium. This was repeated for ~7 days. Monday to Friday, cells were isolated daily at 27°C and scored for survival every 24 hours. Over the weekend, isolated cells were placed at 18°C and scored for survival after 72 hours. Daily re-isolation was resumed during the following week, until death of all lines was observed in the condensin KDs.

For the immunofluorescence staining of MICs in vegetative cells, *en masse* cultures of *P. tetraurelia* cells were subjected to *SMC4.1* or *ND7* KD for 3 days, with daily dilution to 30 cells/mL into freshly induced RNAi medium. Growth arrest was observed between 1 and 2 days of incubation at 27°C. Cells were collected on the 3$^{rd}$ day and immunostained using rabbit anti-γ-tubulin antibodies (Klotz et al., 2003) and AF 568-conjugated secondary anti-rabbit antibodies (antibodies and working dilutions are listed in Table S4).

Survival plots were generated with R using the following packages: *tidyverse* (2.0.0), *survival* (3.8-3) and *survminer* (0.5.0). The MIC and MAC quantification histograms and the *SMC4.1* KD complementation curves were generated using *gridExtra* (2.3), *reshape2* (1.4.4) and *ggplot2* (3.4.2).

#### Co-immunoprecipitation (Co-IP) from co-injected cells

Vegetative *Paramecium* 51 *nd7-1* mutant cells (Dubois et al., 2017) were micro-injected with a 5:5:1 mixture (4 µg/µl final concentration) of linearized DNA from plasmids p0460, p0464 and pGemND7, carrying the RNAi-resistant *CAPD2.3*-HA* and *SMC4.2*-FLAG* fusion transgenes, and the wildtype *ND7* gene, respectively. Triple transformants were selected according to their ability to discharge trichocysts and the copy number of each fusion transgene, as estimated by qPCR (see Zenodo).

Selected triple transformants were grown in 800-mL cultures and starved to induce autogamy upon endogenous *CAPD2.3* and *SMC4.2* KDs to maximize the amount of each tagged subunit in the developing new MACs. Enriched nuclear preparations were obtained from autogamous cells at T5-T10 (~2.5 x $10^6$ cells) as described (Zangarelli et al., 2022), with a few modifications. The cell lysate was washed once in 5 mL of washing solution B (0.25 M sucrose, 10 mM $MgCl_2$, 10 mM Tris pH 7.4) supplemented with 2× protease inhibitor cocktail set I (PIC-S1; Calbiochem 539131). The nuclei-enriched pellet was collected by centrifugation at 1,000g for 2 min at 4 °C and either processed immediately, or resuspended in washing solution B + 13% glycerol supplemented with 2x PIC-S1 and stored at -80°C.

Protein extraction was performed as described (Miró-Pina et al., 2022) with the following modifications: the nuclear pellet was resuspended in 1 volume (~300 µl) of 2X nuclear extraction buffer B (100 mM Hepes pH 7.8, 100 mM KCl, 300 mM NaCl, 0.2 mM EDTA, 20% glycerol, 2 mM DTT, 0.02% Nonidet P-40, 4x PIC-S1) and incubated overnight at 4°C with shaking. The soluble fraction at 150 mM NaCl was recovered by centrifugation at 10,000g for 10 min at 4°C.

Before IP, primary antibodies were pre-adsorbed on Protein G Sepharose 4 Fast Flow beads (Cytiva, #17061801) as described (Eguether et al., 2014), with the following modifications. The appropriate volume of preswollen bead slurry (60 µL per condition) was washed three times at 4°C in 1X PBS (137 mM NaCl, 2.7 mM KCl, 10 mM $Na_2HPO_4$, 1.8 mM $KH_2PO_4$, pH 7.4) and beads were collected by centrifugation at 4°C for 2 min (280g). For each IP, 30 µl of bead pellet (~60 µl of the initial bead slurry) were resuspended in 250 µl PBS and incubated overnight at 4°C under shaking with 5 µg of mouse monoclonal anti-HA (HA7, H9658, Sigma-Aldrich) or anti-FLAG (M2, F1804, Sigma Aldrich) antibodies, or with Normal Mouse IgG Polyclonal Antibody (#12371, Millipore). Unbound antibodies were removed by washing the beads twice in PBS, then one time in 1x nuclear extraction buffer B.

For IP, antibody-coupled beads were pelleted, resuspended in ~200 µl of soluble fraction and the mixture was incubated for 3 hours at 4°C on a spinning wheel. Beads were washed four times in TEGN buffer (20 mM Tris pH 8, 0.1 mM EDTA, 10% Glycerol, 150 mM NaCl, 0.01% Nonidet P-40), and resuspended in 60 µl 1X Laemmli sample buffer supplemented with β-mercaptoethanol (10% v/v final). For SDS-PAGE electrophoresis and western blot analysis, the bound fraction was separated from the beads by heating at 95°C for 3 min and centrifugation at 12,000 g for 30 sec before loading on a 4-15% polyacrylamide Mini-PROTEAN® TGX™ Precast Protein Gel (BIO-RAD).

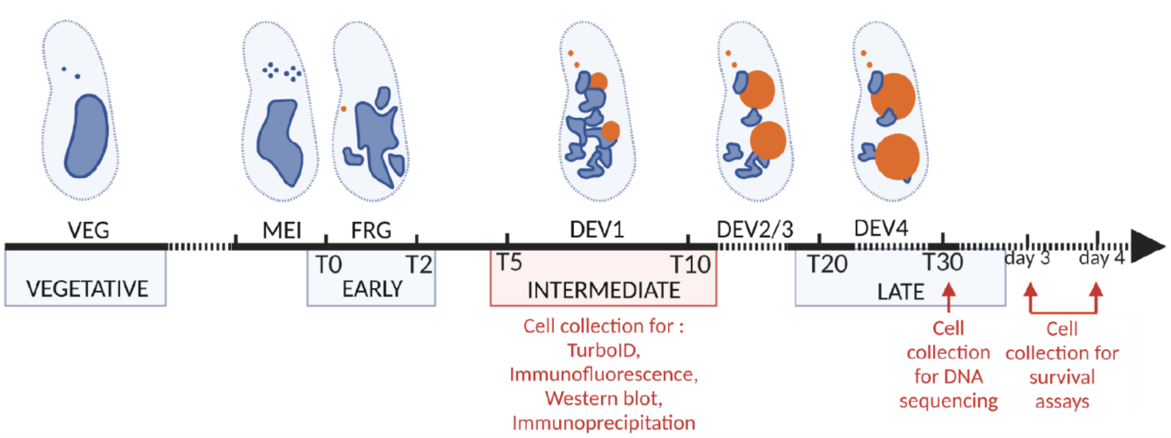


**Supplementary Figure S1. Experimental workflow during autogamy**

Diagram showing the correspondence between autogamy stages (VEG, MEI, FRG, DEV1 to DEV4) and gene expression clusters (VEGETATIVE, EARLY, INTERMEDIATE, LATE) (Arnaiz et al., 2017; Bazin-Gélis et al., 2023). Time-points are in hours, with T0 defined as the time when 50% of cells in the population harbor a fragmented MAC. The progression of nuclear shapes is represented on top (blue: old MICs and MAC; orange: zygotic nucleus, new MICs and MACs). Cell collection time-points for the experiments presented in this study are indicated in red. Days 3 and 4 correspond to ~T50 and ~T72, respectively. Created with BioRender.com

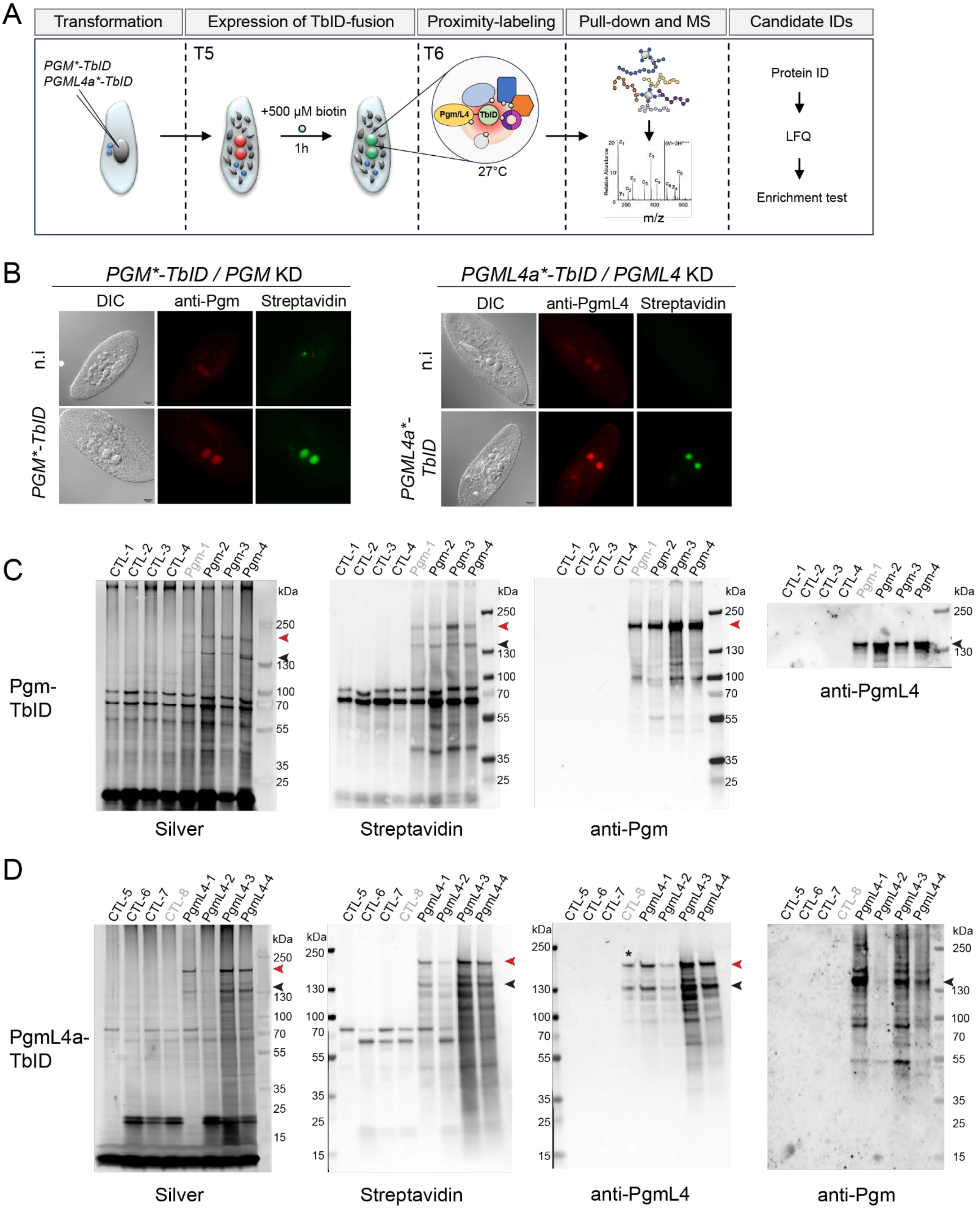


**Supplementary Figure S2. TbID-based proximity labeling in *Paramecium***

**(A)** Overview of the TbID-MS workflow. RNAi-resistant TbID fusion transgenes (*PGM*-TbID* or *PGML4a*-TbID*, carried by plasmids p0380/p0381 or p0398/p0400, respectively) are delivered into the MAC of vegetative cells via microinjection. Non-injected controls and transformed clones are subjected to RNAi against each corresponding endogenous gene. At T5 during autogamy, biotin is added for 1 hour before cell collection at T6. Biotinylated proteins from total cell lysates are captured on streptavidin beads and submitted to mass spectrometry for protein identification and label-free quantification.

**(B)** Subcellular localization of TbID fusions and TbID-induced biotinylated proteins. Pgm-TbID and PgmL4a-TbID were expressed from p0380 and p0400, respectively. n.i: non-injected controls. Immunofluorescence and streptavidin labeling of autogamous cells was performed at T6, followed by epifluorescence imaging. Scale bars: 10 µm

**(C)** Polyacrylamide gel electrophoresis and western blot analysis of proteins bound on streptavidin beads, for non-injected controls (CTL) or cells expressing Pgm-TbID from plasmid p0381. All cultures were subjected to RNAi against endogenous *PGM*. Four replicates are shown for each condition. In grey: samples that were excluded from the statistical analysis of mass spectrometry data. Four gels were run in parallel: one for direct silver staining (Silver), the other three for western blot analysis using streptavidin HRP (Streptavidin) or anti-Pgm or anti-PgmL4 antibodies. Red arrowheads point to the full-length Pgm-TbID fusion, labeled with streptavidin-HRP or anti-Pgm antibodies. Black arrowheads point to endogenous PgmL4, labeled with anti-PgmL4 antibodies.

**(D)** Same as (C) for cells expressing PgmL4a-FLAG-TbID from plasmid p0398, and subjected to *PGML4a+b* RNAi. Red arrowheads point to full-length PgmL4a-TbID, black arrowheads to endogenous Pgm and a proteolytic product of PgmL4a-TbID. The asterisk in the third panel indicates that the PgmL4-1 sample leaked into the CTL-8 lane.

In (B-D), the survival of post-autogamous sexual progeny was examined to confirm that microinjected RNAi-resistant TbID fusion transgenes are functional and complement RNAi-mediated KD of their respective endogenous counterpart (see Supplementary File S1). Survival assays were performed in the absence of biotin for cells expressing PgmL4a-TbID.

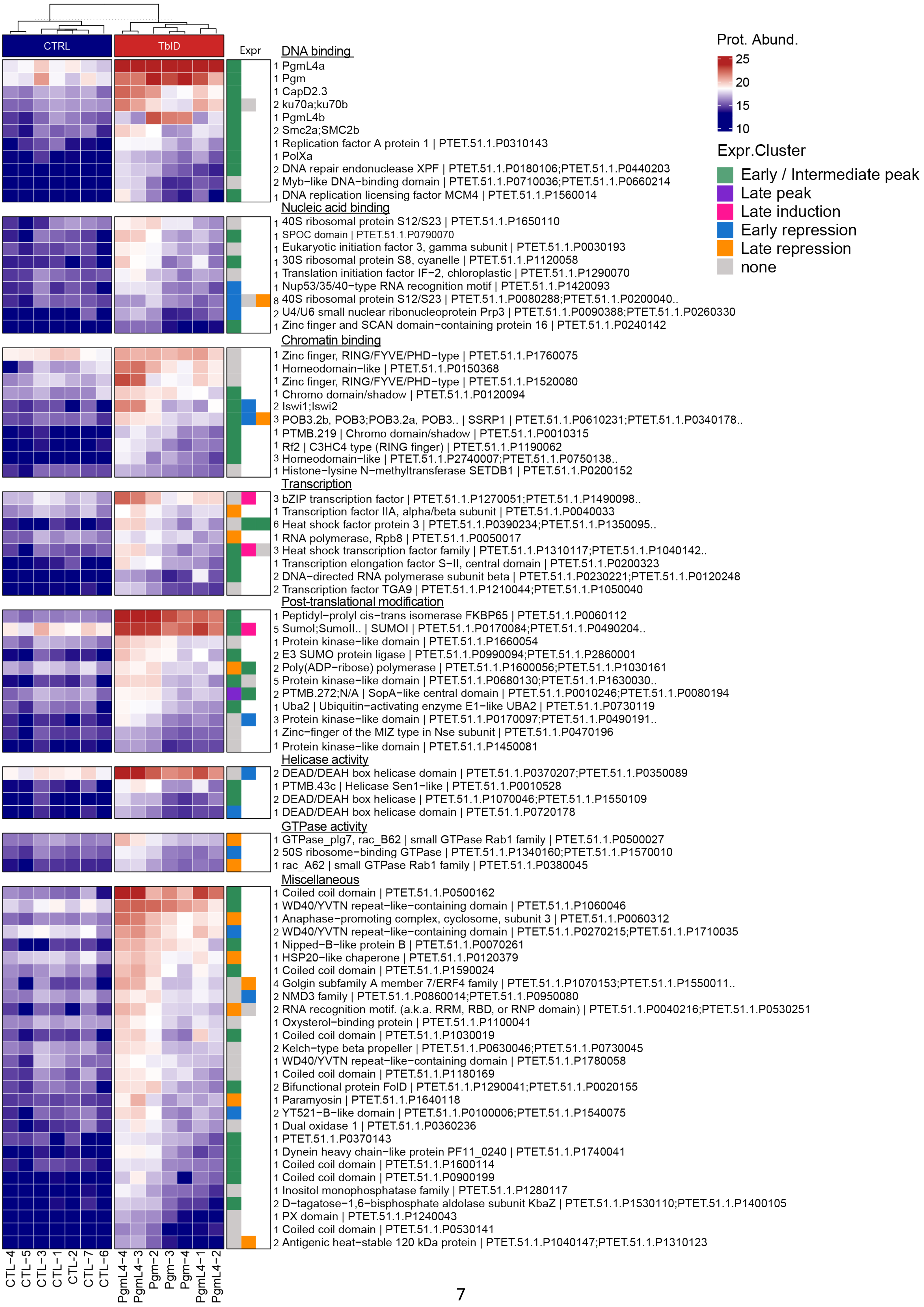

CTRL
TbID
Expr
Prot. Abund.
25
20
15
10
Expr.Cluster
Early / Intermediate peak
Late peak
Late induction
Early repression
Late repression
none
DNA binding
1 PgmL4a
1 Pgm
1 CapD2.3
2 ku70a;ku70b
1 PgmL4b
2 Smc2a;SMC2b
1 Replication factor A protein 1 | PTET.51.1.P0310143
1 PolXa
2 DNA repair endonuclease XPF | PTET.51.1.P0180106;PTET.51.1.P0440203
2 Myb−like DNA−binding domain | PTET.51.1.P0710036;PTET.51.1.P0660214
1 DNA replication licensing factor MCM4 | PTET.51.1.P1560014
Nucleic acid binding
1 40S ribosomal protein S12/S23 | PTET.51.1.P1650110
1 SPOC domain | PTET.51.1.P0790070
1 Eukaryotic initiation factor 3, gamma subunit | PTET.51.1.P0030193
1 30S ribosomal protein S8, cyanelle | PTET.51.1.P1120058
1 Translation initiation factor IF−2, chloroplastic | PTET.51.1.P1290070
1 Nup53/35/40−type RNA recognition motif | PTET.51.1.P1420093
8 40S ribosomal protein S12/S23 | PTET.51.1.P0080288;PTET.51.1.P0200040..
2 U4/U6 small nuclear ribonucleoprotein Prp3 | PTET.51.1.P0090388;PTET.51.1.P0260330
1 Zinc finger and SCAN domain−containing protein 16 | PTET.51.1.P0240142
Chromatin binding
1 Zinc finger, RING/FYVE/PHD−type | PTET.51.1.P1760075
1 Homeodomain−like | PTET.51.1.P0150368
1 Zinc finger, RING/FYVE/PHD−type | PTET.51.1.P1520080
1 Chromo domain/shadow | PTET.51.1.P0120094
2 Iswi1;Iswi2
3 POB3.2b, POB3;POB3.2a, POB3.. | SSRP1 | PTET.51.1.P0610231;PTET.51.1.P0340178..
1 PTMB.219 | Chromo domain/shadow | PTET.51.1.P0010315
1 Rf2 | C3HC4 type (RING finger) | PTET.51.1.P1190062
3 Homeodomain−like | PTET.51.1.P2740007;PTET.51.1.P0750138..
1 Histone−lysine N−methyltransferase SETDB1 | PTET.51.1.P0200152
Transcription
3 bZIP transcription factor | PTET.51.1.P1270051;PTET.51.1.P1490098..
1 Transcription factor IIA, alpha/beta subunit | PTET.51.1.P0040033
6 Heat shock factor protein 3 | PTET.51.1.P0390234;PTET.51.1.P1350095..
1 RNA polymerase, Rpb8 | PTET.51.1.P0050017
3 Heat shock transcription factor family | PTET.51.1.P1310117;PTET.51.1.P1040142..
1 Transcription elongation factor S−II, central domain | PTET.51.1.P0200323
2 DNA−directed RNA polymerase subunit beta | PTET.51.1.P0230221;PTET.51.1.P0120248
2 Transcription factor TGA9 | PTET.51.1.P1210044;PTET.51.1.P1050040
Post-translational modification
1 Peptidyl−prolyl cis−trans isomerase FKBP65 | PTET.51.1.P0060112
5 Sumol;SumolI.. | SUMOI | PTET.51.1.P0170084;PTET.51.1.P0490204..
1 Protein kinase−like domain | PTET.51.1.P1660054
2 E3 SUMO protein ligase | PTET.51.1.P0990094;PTET.51.1.P2860001
2 Poly(ADP−ribose) polymerase | PTET.51.1.P1600056;PTET.51.1.P1030161
5 Protein kinase−like domain | PTET.51.1.P0680130;PTET.51.1.P1630030..
2 PTMB.272;N/A | SopA−like central domain | PTET.51.1.P0010246;PTET.51.1.P0080194
1 Uba2 | Ubiquitin−activating enzyme E1−like UBA2 | PTET.51.1.P0730119
3 Protein kinase−like domain | PTET.51.1.P0170097;PTET.51.1.P0490191..
1 Zinc−finger of the MIZ type in Nse subunit | PTET.51.1.P0470196
1 Protein kinase−like domain | PTET.51.1.P1450081
Helicase activity
2 DEAD/DEAH box helicase domain | PTET.51.1.P0370207;PTET.51.1.P0350089
1 PTMB.43c | Helicase Sen1−like | PTET.51.1.P0010528
2 DEAD/DEAH box helicase | PTET.51.1.P1070046;PTET.51.1.P1550109
1 DEAD/DEAH box helicase domain | PTET.51.1.P0720178
GTPase activity
1 GTPase_plg7, rac_B62 | small GTPase Rab1 family | PTET.51.1.P0500027
2 50S ribosome−binding GTPase | PTET.51.1.P1340160;PTET.51.1.P1570010
1 rac_A62 | small GTPase Rab1 family | PTET.51.1.P0380045
Miscellaneous
1 Coiled coil domain | PTET.51.1.P0500162
1 WD40/YVTN repeat−like−containing domain | PTET.51.1.P1060046
1 Anaphase−promoting complex, cyclosome, subunit 3 | PTET.51.1.P0060312
2 WD40/YVTN repeat−like−containing domain | PTET.51.1.P0270215;PTET.51.1.P1710035
1 Nipped−B−like protein B | PTET.51.1.P0070261
1 HSP20−like chaperone | PTET.51.1.P0120379
1 Coiled coil domain | PTET.51.1.P1590024
4 Golgin subfamily A member 7/ERF4 family | PTET.51.1.P1070153;PTET.51.1.P1550011..
2 NMD3 family | PTET.51.1.P0860014;PTET.51.1.P0950080
2 RNA recognition motif. (a.k.a. RRM, RBD, or RNP domain) | PTET.51.1.P0040216;PTET.51.1.P0530251
1 Oxysterol−binding protein | PTET.51.1.P1100041
1 Coiled coil domain | PTET.51.1.P1030019
2 Kelch−type beta propeller | PTET.51.1.P0630046;PTET.51.1.P0730045
1 WD40/YVTN repeat−like−containing domain | PTET.51.1.P1780058
1 Coiled coil domain | PTET.51.1.P1180169
2 Bifunctional protein FolD | PTET.51.1.P1290041;PTET.51.1.P0020155
1 Paramyosin | PTET.51.1.P1640118
2 YT521−B−like domain | PTET.51.1.P0100006;PTET.51.1.P1540075
1 Dual oxidase 1 | PTET.51.1.P0360236
1 PTET.51.1.P0370143
1 Dynein heavy chain−like protein PF11_0240 | PTET.51.1.P1740041
1 Coiled coil domain | PTET.51.1.P1600114
1 Coiled coil domain | PTET.51.1.P0900199
1 Inositol monophosphatase family | PTET.51.1.P1280117
2 D−tagatose−1,6−bisphosphate aldolase subunit KbaZ | PTET.51.1.P1530110;PTET.51.1.P1400105
1 PX domain | PTET.51.1.P1240043
1 Coiled coil domain | PTET.51.1.P0530141
2 Antigenic heat−stable 120 kDa protein | PTET.51.1.P1040147;PTET.51.1.P1310123
CTL−4
CTL−5
CTL−3
CTL−1
CTL−2
CTL−7
CTL−6
PgmL4−4
PgmL4−3
Pgm−2
Pgm−3
Pgm−4
PgmL4−1
PgmL4−2

**Supplementary Figure S3. Full list of significantly enriched protein groups in both Pgm-TbID and PgmL4-TbID experiments**

Heatmap shows protein abundance in each sample for the complete list of protein groups shared between the two sets of experiments. Left (CTL): non-injected controls; middle (TbID): TbID samples. Each line corresponds to one protein group. Groups are categorized according to putative molecular function. Within each category, groups are ranked in descending order of mean protein abundance, calculated as the arithmetic mean across TbID samples from the PgmL4-TbID and Pgm-TbID experiments. Right panel (Expr): color-coded expression profile(s) of the encoding genes from each protein group (Arnaiz et al., 2017). The number of proteins in each group is indicated on the right. Group annotation: alias | description | protein accession number. For known PDE proteins, only the alias is shown.

Protein abundance values (normalized, log-transformed abundance) shown in the heatmap are found in columns N to AA of the “TbID_heatmap_full.xlsx” file in Zenodo. For color mapping [blue, white, red], they were linearly interpolated between [13.10, 17.93, 22.76]. Mean protein abundance values (in the TbID samples) are listed under "rank_AREA" in the above file.

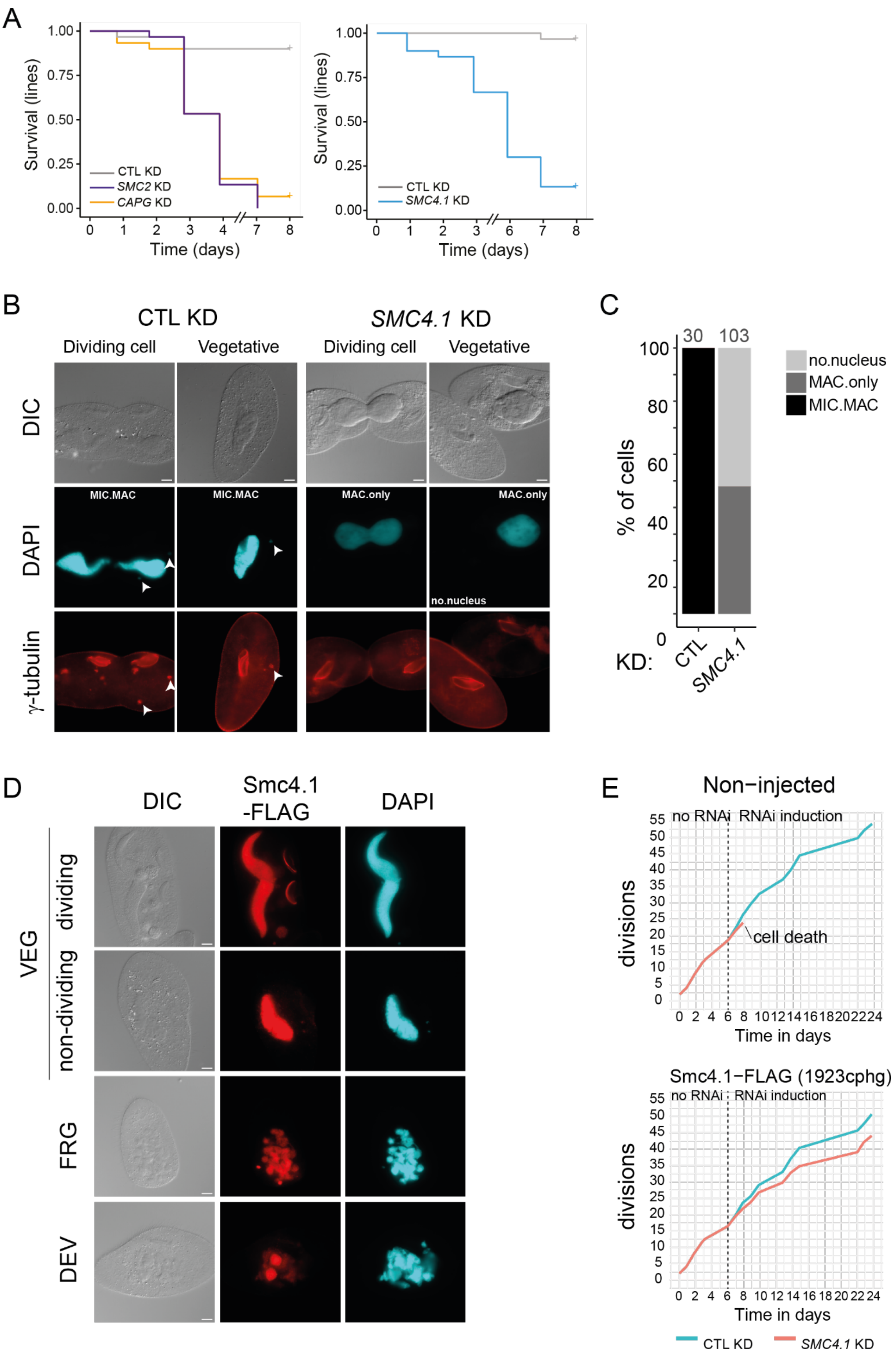


**Supplementary Figure S4: Vegetative phenotypes caused by *SMC2a+b*, *SMC4.1* or *CAPG* KD**

**(A)** Survival analysis of vegetative cell lines subjected to control *ND7* KD (CTL) and to *SMC2*, *CAPG* (left), or S*MC4.1* KD (right). For each condition, 30 cell lines were independently propagated with daily cell isolation in fresh RNAi-inducing medium. The proportions of surviving lines were calculated using the Kaplan-Meier estimator.
**(B)** Immunostaining of γ-tubulin in representative dividing and vegetative cells from *en masse* cultures subjected to *ND7* KD (CTL) and *SMC4.1* KD for three days. Nuclei were stained with DAPI and MICs are labeled with arrowheads. Note that anti γ-tubulin antibodies label the cortex in addition to the MICs (Klotz et al., 2003). Scale bar: 10 µm.
**(C)** Quantification of the number of cells with at least one MIC and one MAC (MIC.MAC), one MAC but no visible MIC (MAC.only), or without any nucleus (no.nucleus), in the experiment shown in (B). The total number of cells that were examined is indicated above each bar.
**(D)** Nuclear localization of Smc4.1-FLAG during vegetative growth (VEG) and autogamy (FRG and DEV stages, see Figure S1), in cells harboring a *SMC4.1-FLAG* transgene and subjected to *ND7* control KD. Scale bar: 10 µm.
**(E)** Complementation of *SMC4.1* KD during vegetative growth by the transgene used in D. Control non-injected cells (top) and cells harboring the *SMC4.1-FLAG* transgene (bottom) (cphg: transgene copy number per haploid genome) were transferred to *SMC4.1* RNAi-inducing medium at day 6 (dotted line) and maintained in vegetative growth for 24 days by daily transfer to fresh RNAi medium, as in (A). The curves show the cumulative count of vegetative divisions under each condition.

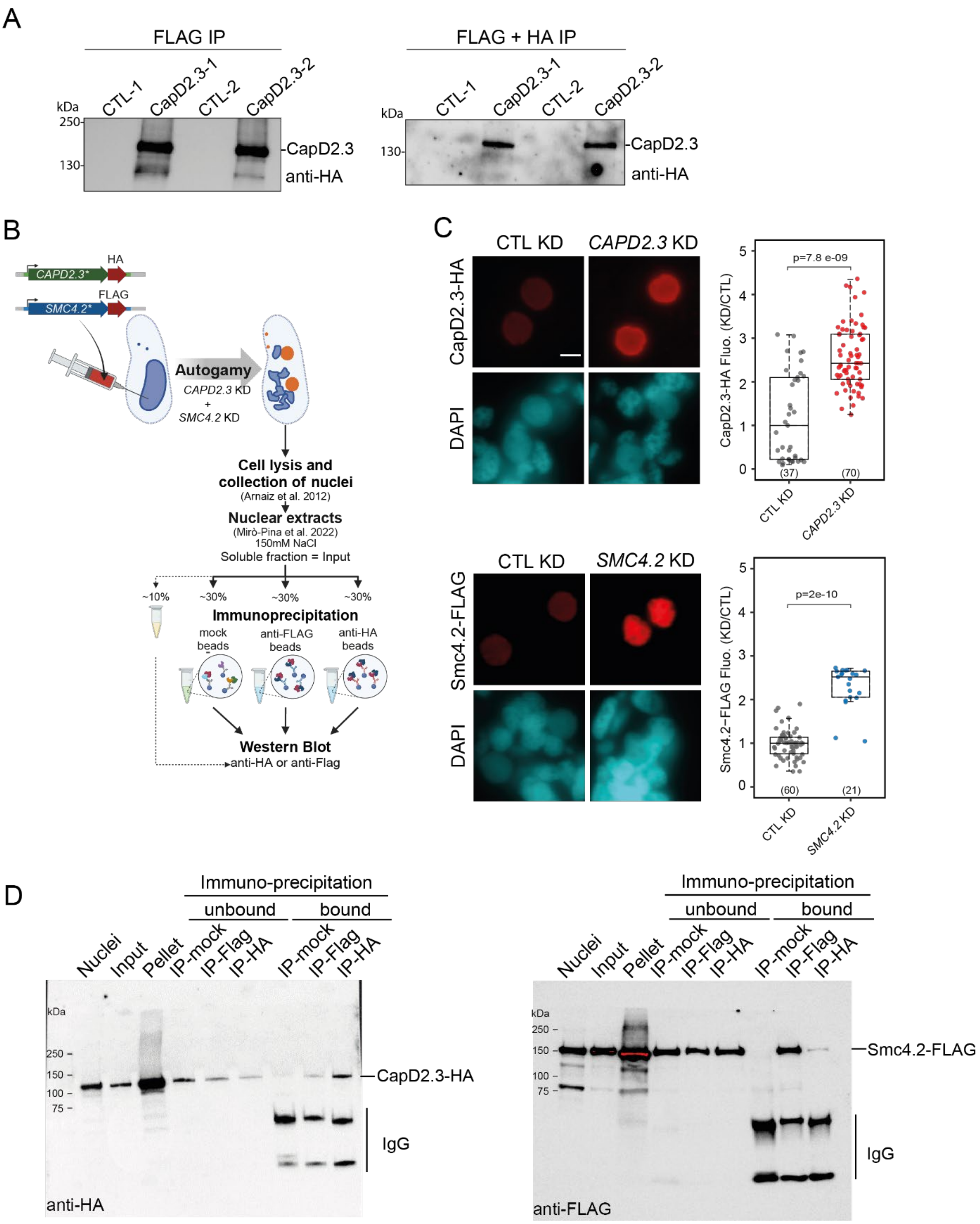


**Supplementary Figure S5. *In vivo* association of CapD2.3 and Smc4.2**

**(A)** CapD2.3 tandem immunoprecipitation replicates of the experiment shown in Figure 2B-C. FLAG (left) and FLAG+HA (right) immunoprecipitations were performed from nuclear extracts of non-injected cells (CTL-1 and CTL-2) and cells expressing a CapD2.3-3xFLAG-HA functional protein (CapD2.3-1 and CapD2.3-2). The western blot analysis of CapD2.3 after tandem affinity purification (FLAG +HA IP) for replicate CapD2.3-1 is also displayed in Figure 2B. Anti-HA antibodies are used to detect CapD2.3.

**(B)** Experimental procedure for the co-immunoprecipitation of Smc4.2-FLAG and CapD2.3-HA following co-expression in autogamous cells.

**(C)** Immunofluorescence labeling of CapD2.3-HA (left) and Smc4.2-FLAG (right) in the presence or absence of their respective endogenous counterpart. Cells harboring an RNAi-resistant *CAPD2.3-HA* or *SMC4.2-FLAG* transgene were subjected to endogenous *CAPD2.3* or *SMC4.2* KD, respectively, and to *ND7* KD (CTL). Scale bar: 5 µm. The quantification of HA or FLAG fluorescence intensities in developing new MACs under each condition, divided by the median fluorescence intensity in their respective control, is shown on the right of each image panel. Sample differences compared to CTL KDs were tested using a Mann-Whitney U test. Quantification boxplots were generated using the following R packages: tidyverse (2.0.0), ggsci (3.2.0) and ggpubr (0.6.0). The number of imaged nuclei is indicated below each box.

**(D)** Western blot analysis of the immuno-precipitates (IP) obtained using mouse IgG (mock), anti-FLAG or anti-HA antibodies, from the nuclear extracts of cells co-expressing Smc4.2-FLAG and CapD2.3-HA. The antibodies used to reveal immunoprecipitated proteins are indicated at the bottom left of each blot.

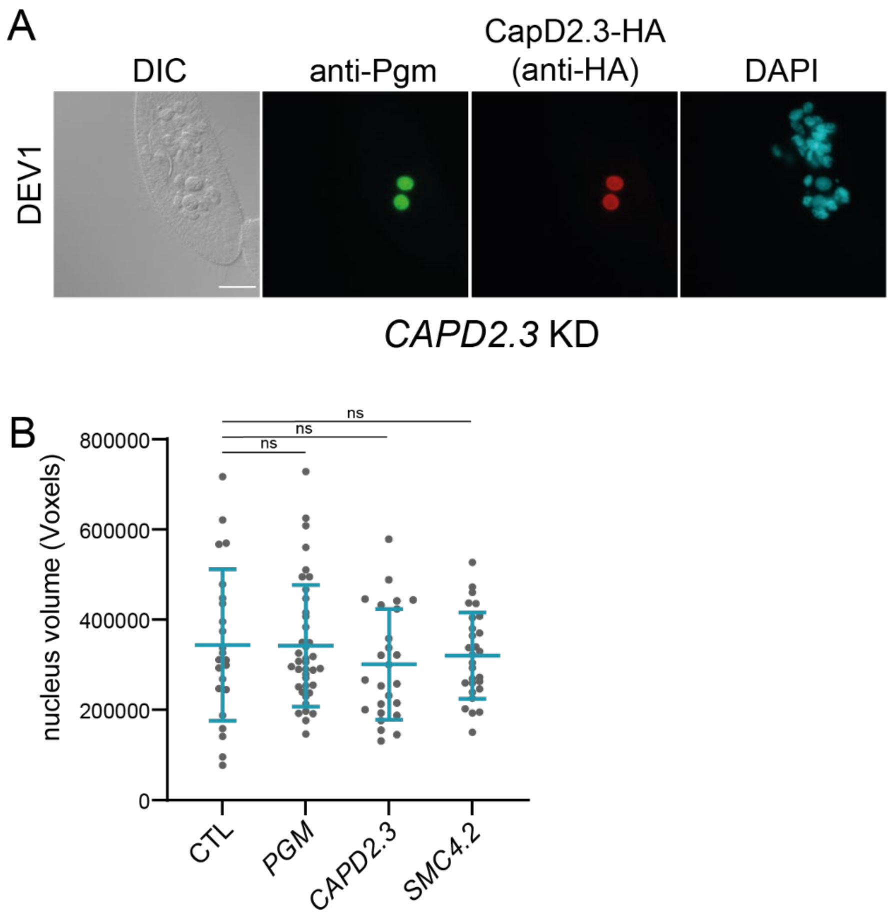


**Supplementary Figure S6: Controls of the localization experiments shown in Figure 3**
**(A)** Immunofluorescence staining of Pgm and CapD2.3-HA at T5-T10 (DEV1, see Figure S1) in a cell expressing the RNAi-resistant *CAPD2.3-HA* transgene and subjected to *CAPD2.3* KD (epifluorescence microscopy images). New developing MACs are labeled with anti-Pgm antibodies, CapD2.3-HA is detected using anti-HA antibodies. DIC: Differential interference contrast. Scale bar: 10 µm.
**(B)** Dot plot of estimated developing new MAC volume in voxels from the same data as in Figure 3F. Estimation of nuclear volume indicated comparable developmental stages in the different conditions. Bars correspond to mean ± SD. Mann-Whitney statistical test. n.s: non-significant.

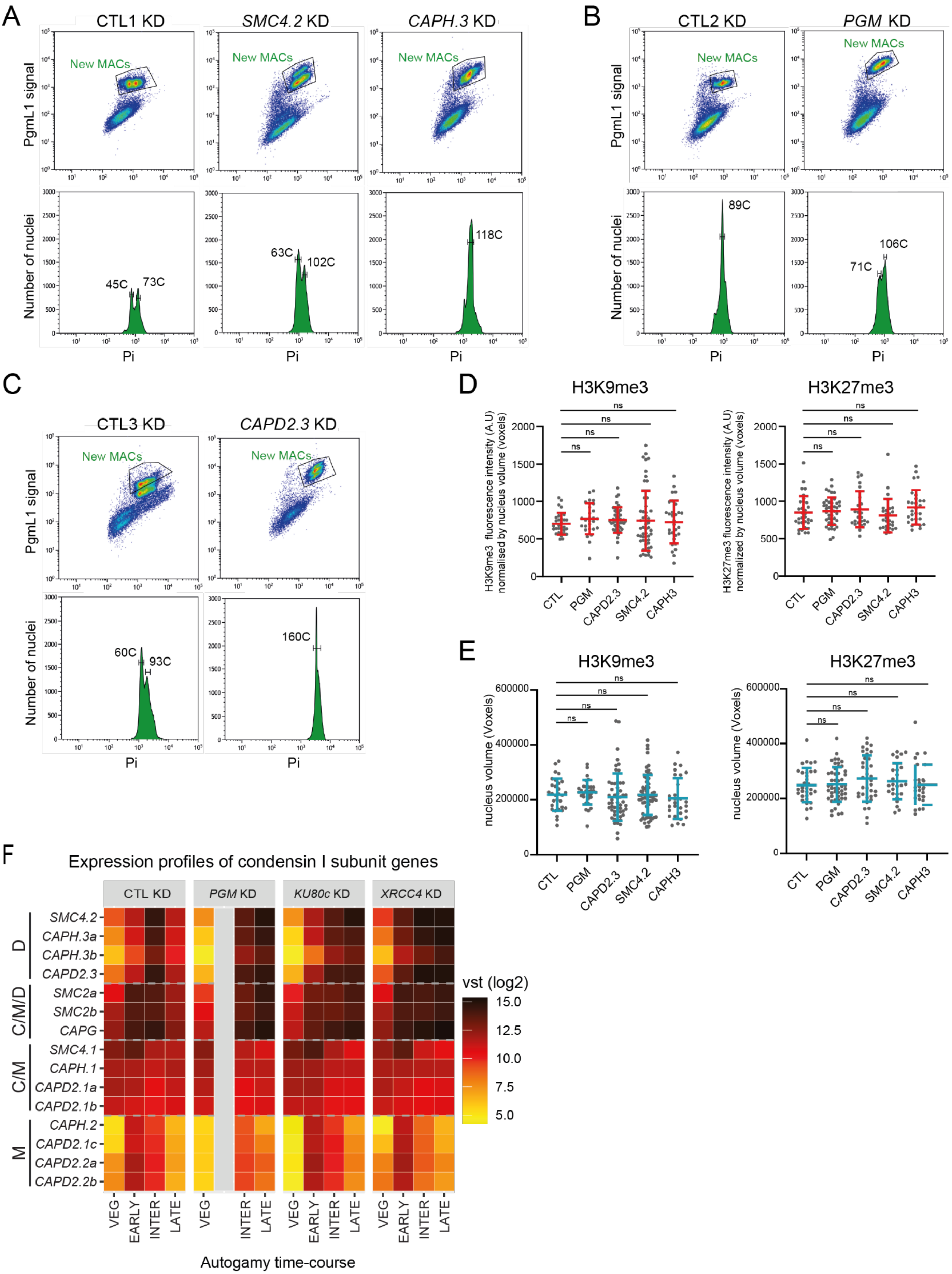
A
CTL1 KD
SMC4.2 KD
CAPH.3 KD
New MACs
PgmL1 signal
Number of nuclei
45C
73C
63C
102C
118C
Pi
B
CTL2 KD
PGM KD
89C
71C
106C
C
CTL3 KD
CAPD2.3 KD
60C
93C
160C
D
H3K9me3
H3K27me3
H3K9me3 fluorescence intensity (A.U) normalised by nucleus volume (voxels)
H3K27me3 fluorescence intensity (A.U) normalized by nucleus volume (voxels)
ns
CTL
PGM
CAPD2.3
SMC4.2
CAPH3
E
nucleus volume (Voxels)
F
Expression profiles of condensin I subunit genes
CTL KD
PGM KD
KU80c KD
XRCC4 KD
D
C/M/D
C/M
M
SMC4.2
CAPH.3a
CAPH.3b
CAPD2.3
SMC2a
SMC2b
CAPG
SMC4.1
CAPH.1
CAPD2.1a
CAPD2.1b
CAPH.2
CAPD2.1c
CAPD2.2a
CAPD2.2b
VEG
EARLY
INTER
LATE
vst (log2)
15.0
12.5
10.0
7.5
5.0
Autogamy time-course

**Supplementary Figure S7. Controls for the study of programmed DNA elimination and H3 modifications in condensin I-depleted cells**

**(A-C)** Flow cytometry sorting of PgmL1-immunostained nuclei from autogamous control cells (CTL1-3) fed with bacteria containing the empty L4440 vector, and from cells subjected to *SMC4.2*, *CAPH.3* (A), *PGM* (B) or *CAPD2.3* KD (C). Nuclei were collected at T28-T30. On the top graphs, the “new MACs” gate was used for sorting. The DNA content of sorted nuclei was estimated using the histograms below.

**(D)** Quantification of H3K9me3 (left) and H3K27me3 (right) fluorescence intensity in the developing MAC from the same data as in Figure 4C-D. The fluorescence intensity is normalized by the nucleus size (in voxels). Bars correspond to mean ± SD. Mann-Whitney statistical test. n.s: non-significant

**(E)** Dot plot of estimated nucleus (developing MAC) volume in voxels from the same data as in Figure 4C-D. Estimation of nuclear volume indicated comparable developmental stages in the different conditions. Bars correspond to mean ± SD. Mann-Whitney statistical test. n.s: non-significant.

**(F)** Transcriptome analysis of all genes encoding condensin subunits, in control conditions (CTL) and in *PGM*, *KU80c* or *XRCC4* KD (Bazin-Gélis et al., 2023). Expression levels ($log_2$ scale) were normalized using variance stabilizing transformation (vst). Genes are ordered using hierarchical clustering (Pearson correlation) and attributed to the putative condensin I complexes defined in Figure 1B-C (C: constitutive, M: meiotic; D: developmental). Autogamy stages are described in Figure S1.

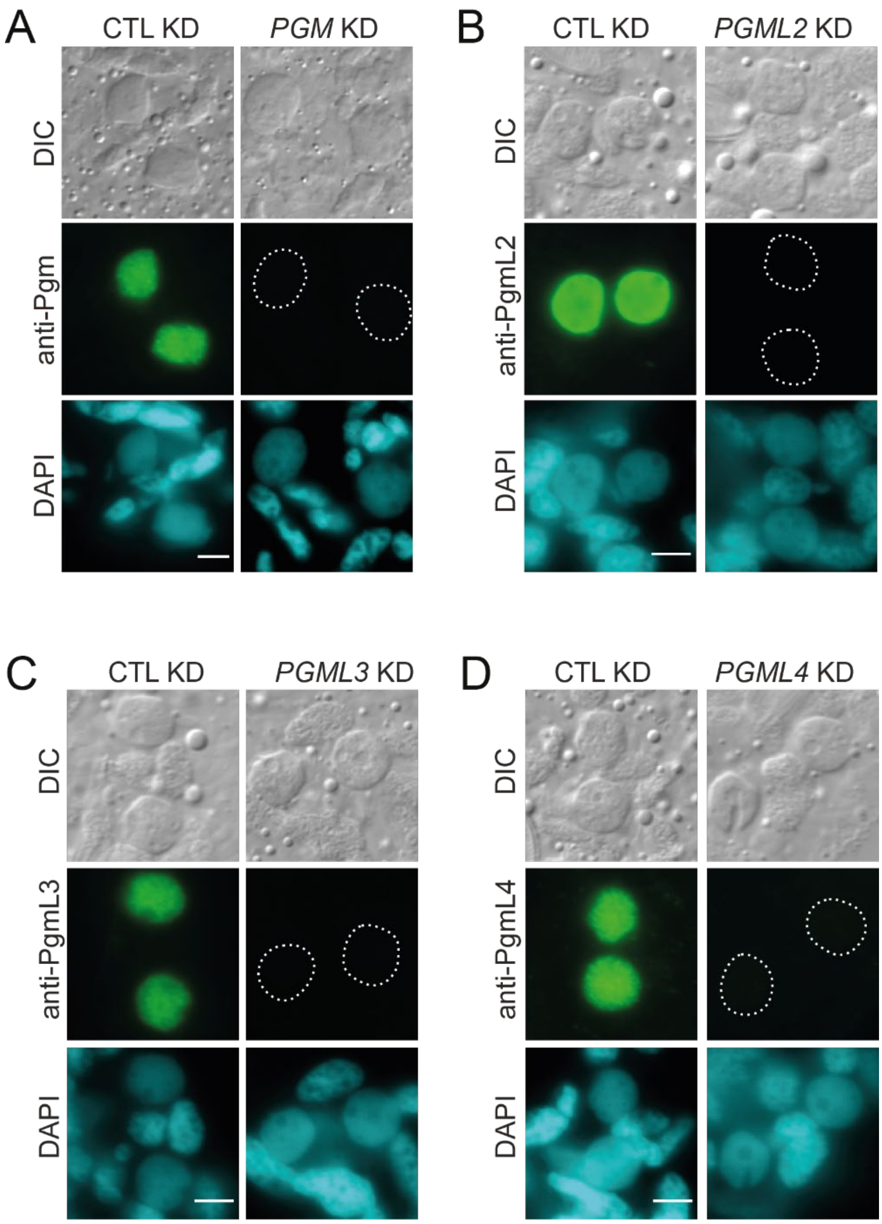


**Supplementary Figure S8. Specificity of anti-Pgm and anti-PgmL antibodies for immunofluorescence labeling of whole cells**

Validation of the specificity of anti-Pgm (GP3) (A), anti-PgmL2 (B), anti-PgmL3a (C) and anti-PgmL4a (D) antibodies by immunofluorescence labelling of fixed autogamous cells collected at T5. In all panels, the control KD (CTL) targeted the *ND7* gene (A) or was performed by feeding cells with bacteria containing the empty L4440 vector (B-D). *PGM* (A) and *PGML2* (B) KDs were single-gene KDs. *PGML3* (C) and *PGML4* (D) KDs targeted two recently duplicated a and b paralogs for each *PGML* family (Bischerour et al., 2018). Scale bar: 5 µm.

Between 15 to 30 cells were examined for each condition. Representative images of cells are shown, with new MACs at the same developmental stage for each KD and its respective CTL, as estimated by size (range 15-53 $\mu m^2$ for A and 30-40 $\mu m^2$ for B-D).

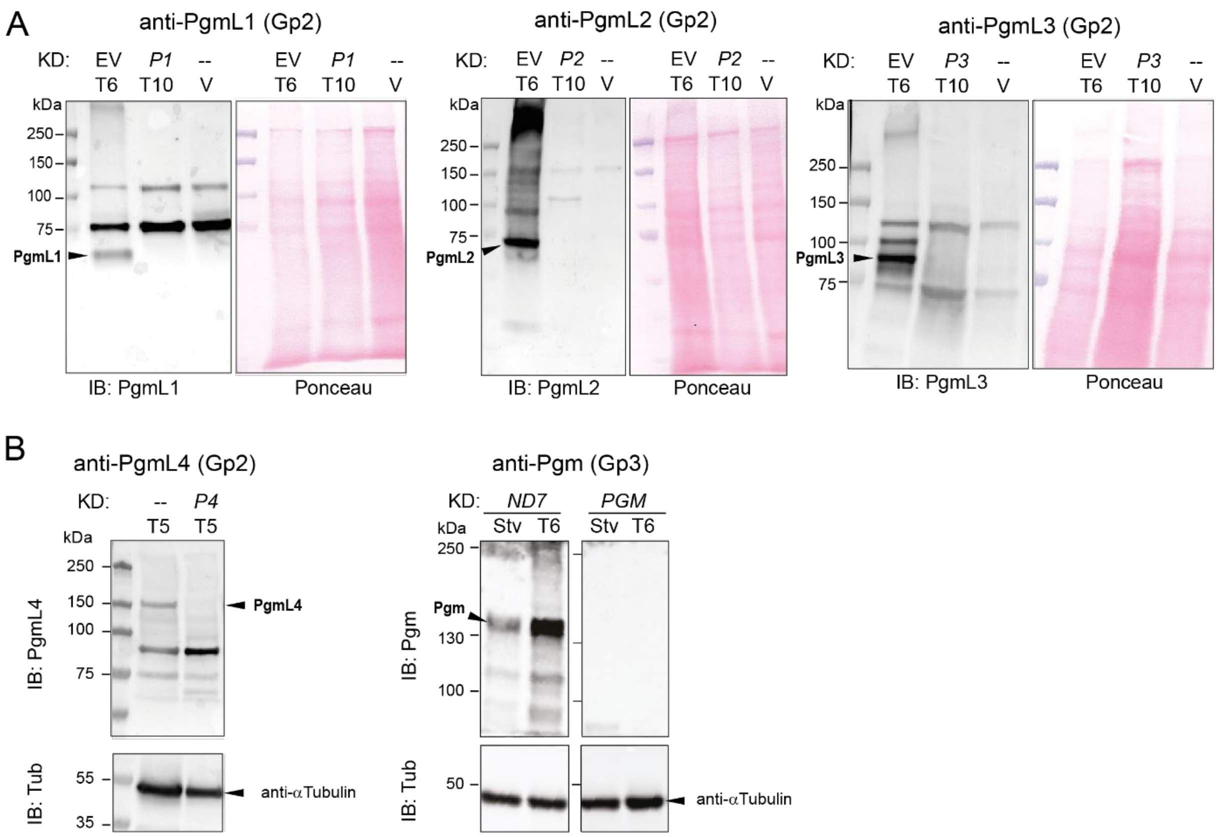


**Supplementary Figure S9. Specificity of anti-Pgm and anti-PgmL antibodies on western blots**

**(A)** Total proteins from autogamous cultures of control cells (EV: empty vector) or cells subjected to *PGML1* (*P1*), *PGML2* (*P2*) or *PGML3* (*P3*) KD (autogamy time-points indicated on top) were tested using anti-PgmL1, PgmL2 and PgmL3 antibodies, respectively. V: vegetative cells grown in standard growth medium (--: no RNAi). A Ponceau-stained loading control is shown on the right of each immunoblot (IB).

**(B)** (Left) Test of anti-PgmL4 antibodies on total proteins from autogamous cells at T5, grown in standard growth medium (--: no RNAi) or upon *PGML4* KD (*P4*). (Right) Test of anti-Pgm antibodies on total proteins from starved (Stv) or autogamous cells at T6, subjected to control (*ND7*) or *PGM* KD. Alpha-tubulin loading controls are shown at the bottom. The full membrane can be found in Zenodo.

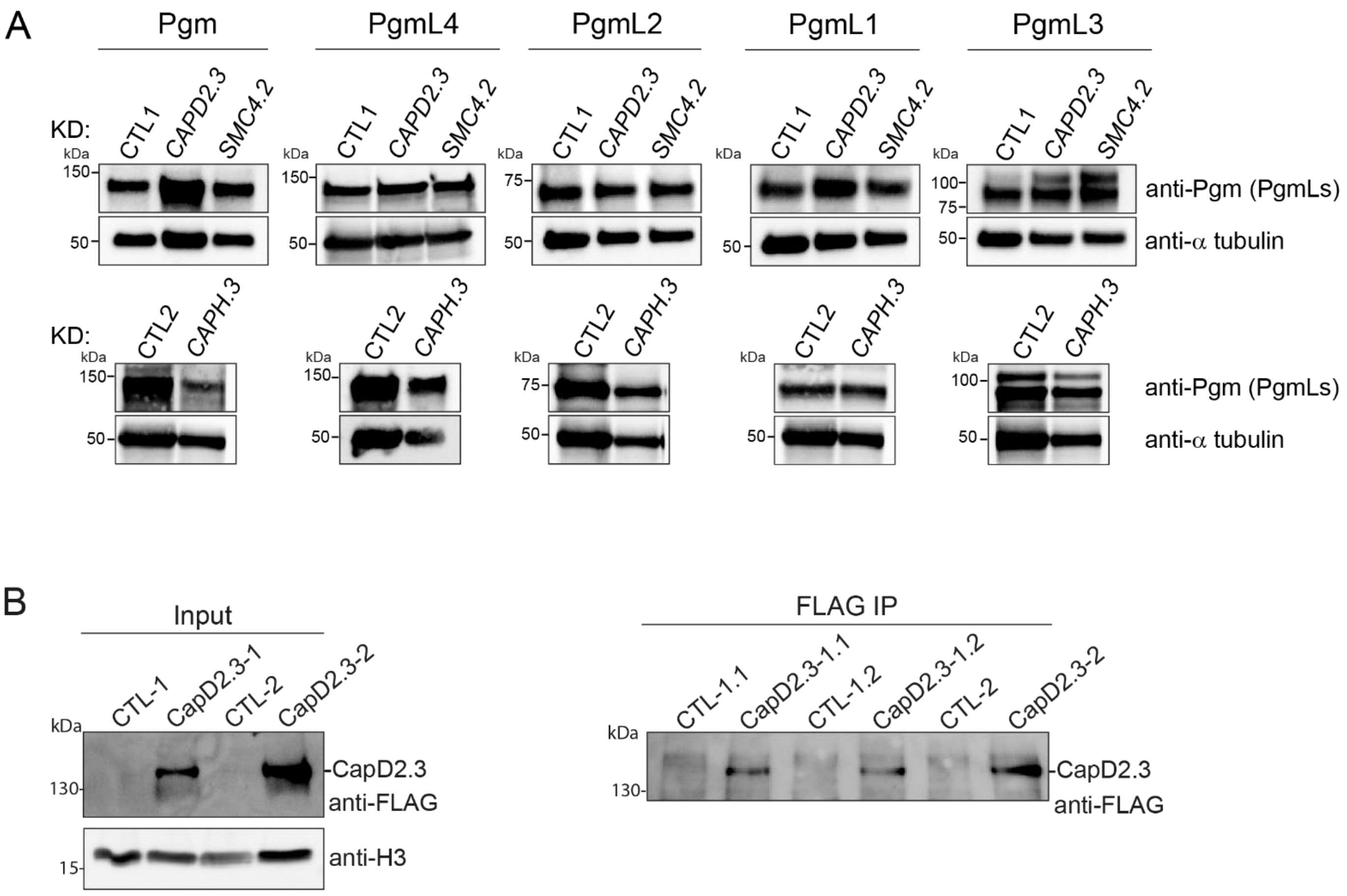


**Supplementary Figure S10. Control western blots for the experiments displayed in Figure 5**

**(A)** Control for Figure 5A. Western blot analysis of Pgm, PgmL4a/b (PgmL4), PgmL2, PgmL1 and PgmL3a/b (PgmL3) in cells subjected to control (CTL1 or CTL2: Empty vector), *CAPD2.3*, *SMC4.2* (top panels) or *CAPH.3* KDs (bottom panels), using the specific antibodies validated in Figure S9. The anti-PgmL5 antibodies do not give a specific signal on western blots of total cellular extracts, which prevented us from controlling PgmL5 expression levels.

**(B)** CapD2.3 immunoprecipitation replicates of the experiment shown in Figure 5B-D. FLAG immunoprecipitation was performed from nuclear extracts of non-injected cells (CTL-1 and CTL-2) and cells expressing a CapD2.3-FLAG-HA functional protein (CapD2.3-1 and CapD2.3-2). Western blot analysis of CapD2.3 before (input) or after affinity purification (FLAG IP). CapD2.3-1.1 and CapD2.3-1.2 are two independent IP replicates from CapD2.3-1 cells. The CapD2.3-2 replicate is displayed in Figure 5C. Anti-FLAG antibodies are used to detect CapD2.3, anti-H3 antibodies are for normalization of the input.

**REFERENCES**


Arnaiz O, Van Dijk E, Bétermier M, Lhuillier-Akakpo M, de Vanssay A, Duharcourt S, Sallet E, Gouzy J, Sperling L. 2017. Improved methods and resources for paramecium genomics: transcription units, gene annotation and gene expression. *BMC Genomics* **18**:483. doi:10.1186/s12864-017-3887-z

Bazin-Gélis M, Eleftheriou E, Zangarelli C, Lelandais G, Sperling L, Arnaiz O, Bétermier M. 2023. Inter-generational nuclear crosstalk links the control of gene expression to programmed genome rearrangement during the Paramecium sexual cycle. *Nucleic Acids Res* **51**:12337–12351. doi:10.1093/nar/gkad1006

Bischerour J, Bhullar S, Denby Wilkes C, Régnier V, Mathy N, Dubois E, Singh A, Swart E, Arnaiz O, Sperling L, Nowacki M, Bétermier M. 2018. Six domesticated PiggyBac transposases together carry out programmed DNA elimination in Paramecium. *Elife* **7**:e37927. doi:10.7554/eLife.37927

Dubois E, Mathy N, Regnier V, Bischerour J, Baudry C, Trouslard R, Betermier M. 2017. Multimerization properties of PiggyMac, a domesticated piggyBac transposase involved in programmed genome rearrangements. *Nucleic acids research* **45**:3204–3216. doi:10.1093/nar/gkw1359

Eguether T, Ermolaeva MA, Zhao Y, Bonnet MC, Jain A, Pasparakis M, Courtois G, Tassin A-M. 2014. The deubiquitinating enzyme CYLD controls apical docking of basal bodies in ciliated epithelial cells. *Nat Commun* **5**:4585. doi:10.1038/ncomms5585

Klotz C, Rulz F, Garreau de Loubresse N, Wright M, Dupuid-Williams P, Beisson J. 2003. Gamma-Tubulin and MTOCs in *Paramecium*. *Protist* **154**:193–209. doi:10.1078/143446103322166509

Miró-Pina C, Charmant O, Kawaguchi T, Holoch D, Michaud A, Cohen I, Humbert A, Jaszczyszyn Y, Chevreux G, Del Maestro L, Ait-Si-Ali S, Arnaiz O, Margueron R, Duharcourt S. 2022. Paramecium Polycomb repressive complex 2 physically interacts with the small RNA-binding PIWI protein to repress transposable elements. *Dev Cell* **57**:1037-1052.e8. doi:10.1016/j.devcel.2022.03.014

Zangarelli C, Arnaiz O, Bourge M, Gorrichon K, Jaszczyszyn Y, Mathy N, Escoriza L, Bétermier M, Régnier V. 2022. Developmental timing of programmed DNA elimination in *Paramecium tetraurelia* recapitulates germline transposon evolutionary dynamics. *Genome Research* **32**:2028–2042. doi:10.1101/gr.277027.122
